# Neuronal timescales are functionally dynamic and shaped by cortical microarchitecture

**DOI:** 10.1101/2020.05.25.115378

**Authors:** Richard Gao, Ruud L. van den Brink, Thomas Pfeffer, Bradley Voytek

## Abstract

Complex cognitive functions such as working memory and decision-making require information maintenance over many timescales, from transient sensory stimuli to long-term contextual cues. While theoretical accounts predict the emergence of a corresponding hierarchy of neuronal timescales, direct electrophysiological evidence across the human cortex is lacking. Here, we infer neuronal timescales from invasive intracranial recordings. Timescales increase along the principal sensorimotor-to-association axis across the entire human cortex, and scale with single-unit timescales within macaques. Cortex-wide transcriptomic analysis shows direct alignment between timescales and expression of excitation- and inhibition-related genes, as well as genes specific to voltage-gated transmembrane ion transporters. Finally, neuronal timescales are functionally dynamic: prefrontal cortex timescales expand during working memory maintenance and predict individual performance, while cortex-wide timescales compress with aging. Thus, neuronal timescales follow cytoarchitectonic gradients across the human cortex, and are relevant for cognition in both short- and long-terms, bridging microcircuit physiology with macroscale dynamics and behavior.

## Introduction

Human brain regions are broadly specialized for different aspects of behavior and cognition. For example, primary sensory neurons are tightly coupled to changes in the environment, firing rapidly to the onset and removal of a stimulus, and showing characteristically short intrinsic timescales (Ogawa and Komatsu, 2010; Runyan et al., 2017). In contrast, neurons in cortical association (or transmodal) regions, such as the prefrontal cortex (PFC), can sustain their activity for many seconds when a person is engaged in working memory (Zylberberg and Strowbridge, 2017), decision-making (Gold and Shadlen, 2007), and hierarchical reasoning (Sarafyazd and Jazayeri, 2019). This persistent activity in the absence of immediate sensory stimuli reflects longer neuronal timescales, which is thought to result from neural attractor states (Wang, 2002; Wimmer et al., 2014) shaped by NMDA-mediated recurrent excitation and fast feedback inhibition (Wang, 1999, 2008), with contributions from other synaptic and cell-intrinsic properties (Duarte and Morrison, 2019; Gjorgjieva et al., 2016).

Recent studies have shown that variations in many such microarchitectural features follow continuous and coinciding gradients along a sensory-to-association axis across the cortex, including cortical thickness, cell density, and distribution of excitatory and inhibitory neurons (Hilgetag and Goulas, 2020; Huntenburg et al., 2018; Wang, 2020). In particular, grey matter myelination (Glasser and Van Essen, 2011), which indexes anatomical hierarchy, varies with the expression of numerous genes related to microcircuit function, such as NMDA receptor and inhibitory cell-type marker genes (Burt et al., 2018). Functionally, specialization of the human cortex, as well as structural and functional connectivity (Margulies et al., 2016), also follow similar macroscopic gradients. In addition to the broad differentiation between sensory and association cortices, there is evidence for a finer hierarchical organization within the frontal cortex (Sarafyazd and Jazayeri, 2019). For example, the anterior-most parts of the PFC are responsible for long timescale goal-planning behavior (Badre and D’Esposito, 2009; Voytek et al., 2015a), while healthy aging is associated with a shift in these gradients such that older adults become more reliant on higher-level association regions to compensate for altered lower-level cortical functioning (Davis et al., 2008).

Despite convergent observations of continuous cortical gradients in structural features and cognitive specialization, there is no direct evidence for a similar gradient of neuronal timescales across the human cortex. Such a gradient of neuronal dynamics is predicted to be a natural consequence of macroscopic variations in synaptic connectivity and microarchitectural features (Chaudhuri et al., 2015; Duarte et al., 2017; Huntenburg et al., 2018; Wang, 2020), and would be a primary candidate for how functional specialization emerges as a result of hierarchical temporal processing (Kiebel et al., 2008). Single-unit recordings in rodents and non-human primates hint at a hierarchy of timescales that increase, or expand, progressively along a posterior-to-anterior axis (Murray et al., 2014; Runyan et al., 2017; Wasmuht et al., 2018), while intracranial recordings and functional neuroimaging data collected during perceptual and cognitive tasks suggest likewise in humans (Baldassano et al., 2017; Honey et al., 2012; Lerner et al., 2011; Watanabe et al., 2019). However, these data are either sparsely sampled across the cortex or do not measure neuronal activity at the cellular and synaptic level directly, prohibiting the full construction of an electrophysiological timescale gradient across the human cortex. As a result, while whole-cortex data of transcriptomic and anatomical variations exist, we cannot take advantage of them to dissect the contributions of synaptic, cellular, and circuit connectivity in shaping fast neuronal timescales, nor ask whether regional timescales are dynamic and relevant for human cognition (Fig. 1A).

**Fig. 1.**
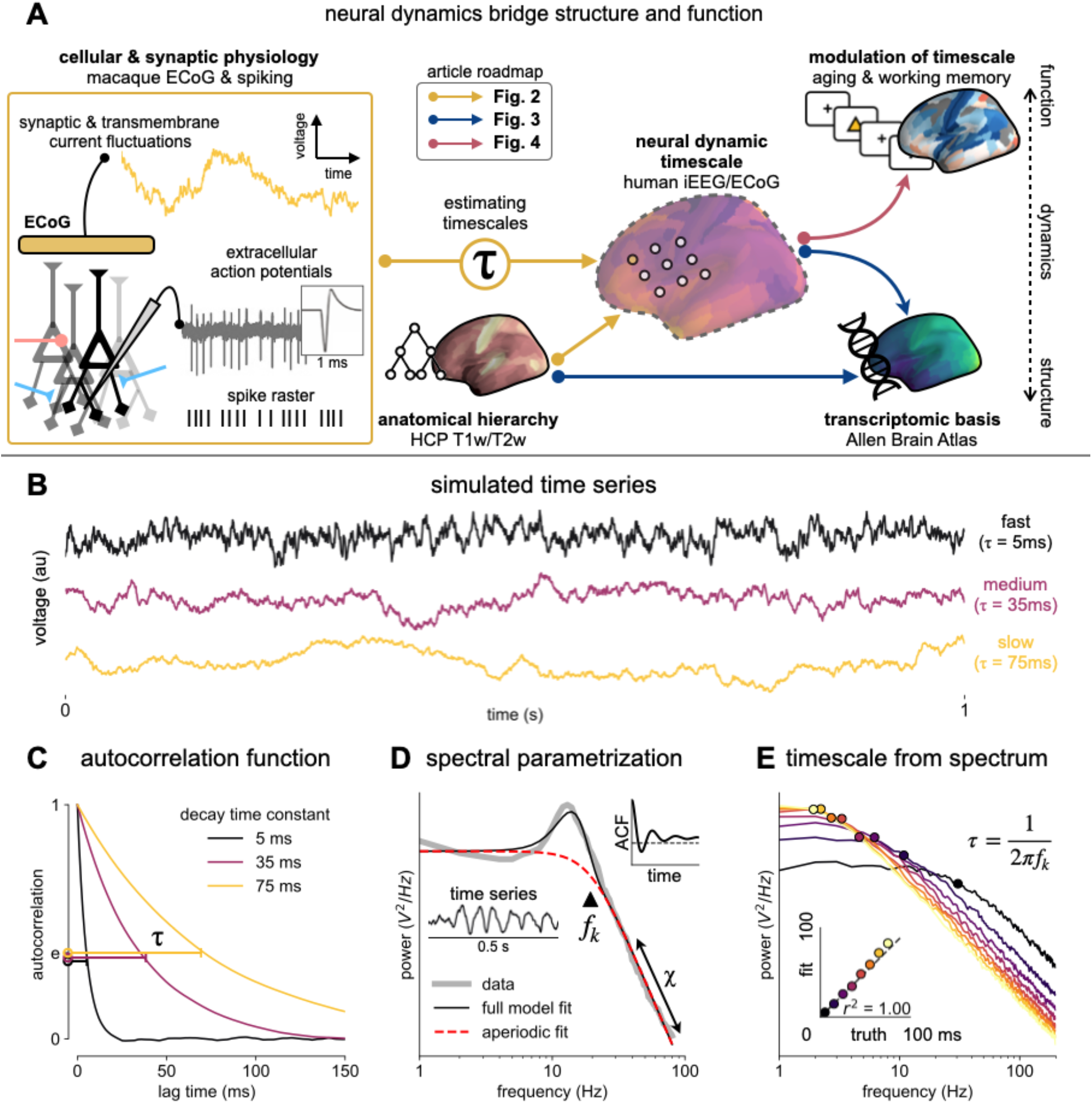
Schematic of study and timescale inference technique. **(A)** in this study, we infer neuronal timescales from intracranial field potential recordings, which reflect integrated synaptic and transmembrane current fluctuations over large neural populations(Buzsáki et al., 2012). Combining multiple open-access datasets (Table S1), we link timescales to known human anatomical hierarchy, dissect its cellular and physiological basis via transcriptomic analysis, and demonstrate its functional modulation during behavior and through aging. **(B)** simulated time series, and their **(C)** autocorrelation functions (ACF), with increasing timescales (decay time constant). **(D)** example ECoG power spectral density (PSD) showing that in frequency domain, timescale is equivalent to the frequency of aperiodic power drop-off (*f_k_*, triangle; insets: time series and ACF). **(E)** accurate extraction of timescale parameters from PSDs of simulated time series in **(B)**.

## Results

### Neuronal timescale can be inferred from frequency domain

To overcome these limitations, we develop a novel computational method for inferring the timescale of neuronal transmembrane current fluctuations from human intracranial electrocorticography (ECoG) recordings (Fig. 1A, box). Neural time series exhibit variable temporal autocorrelation, or timescales, where future values are partially predictable from past values, and predictability decreases with increasing time lags. To demonstrate the effect of varying autocorrelation, we simulate the aperiodic (non-rhythmic) component of neural field potential recordings by convolving Poisson population spikes with exponentially-decaying synaptic kernels (Fig. 1B). Consistent with previous studies, “neuronal timescale” here is defined as the exponential decay time constant (*τ*) of the autocorrelation function (ACF) (Murray et al., 2014)—the time it takes for the ACF to decrease by a factor of *e* (Fig. 1C). Equivalently, we can estimate neuronal timescale from the “characteristic frequency” (*f_k_*) of the power spectral density (PSD), especially when the presence of variable 1/f (χ) and oscillatory components can bias timescale inference from the ACF in time-domain (Fig. 1D). In this study, we apply spectral parameterization (Haller et al., 2018) to extract timescales from intracranial recordings, which decomposes neural PSDs into a combination of oscillatory and aperiodic components, where timescale is inferred from the latter. We validate this approach on PSDs computed from simulated neural time series and show that the model-fitted timescales closely match their ground-truth values (Fig. 1E).

### Timescales follow anatomical hierarchy and are 10x faster than spiking timescales

Applying this technique, we infer a continuous gradient of neuronal timescales across the human cortex and examine its relationship with anatomical hierarchy. We analyze a large dataset of human intracranial (ECoG) recordings of task-free brain activity from 106 epilepsy patients (MNI-iEEG (Frauscher et al., 2018a), see Fig. S1 for electrode coverage), and compare the ECoG-derived timescale gradient to the average T1w/T2w map from the Human Connectome Project, which captures grey matter myelination and indexes the proportion of feedforward vs. feedback connections between cortical regions, defining an anatomical hierarchy (Burt et al., 2018; Glasser and Van Essen, 2011).

Across the human cortex, timescales of fast electrophysiological dynamics (~10-50 ms) predominantly follow a rostrocaudal gradient (Fig. 2A). Consistent with numerous accounts of a principal cortical axis spanning from primary sensory to association regions (Hilgetag and Goulas, 2020; Margulies et al., 2016; Wang, 2020), timescales are shorter in sensorimotor and early visual areas, and longer in association regions, especially posterior parietal, ventral/medial frontal, and medial temporal cortex. Cortical timescales are negatively correlated with T1w/T2w (Fig. 2B, *ρ* = −0.47, *p* < 0.001; adjusted for spatial autocorrelation, see Materials and Methods and Fig. S2), such that timescales are shorter in more heavily myelinated (lower-level) cortical regions.

**Fig. 2.**
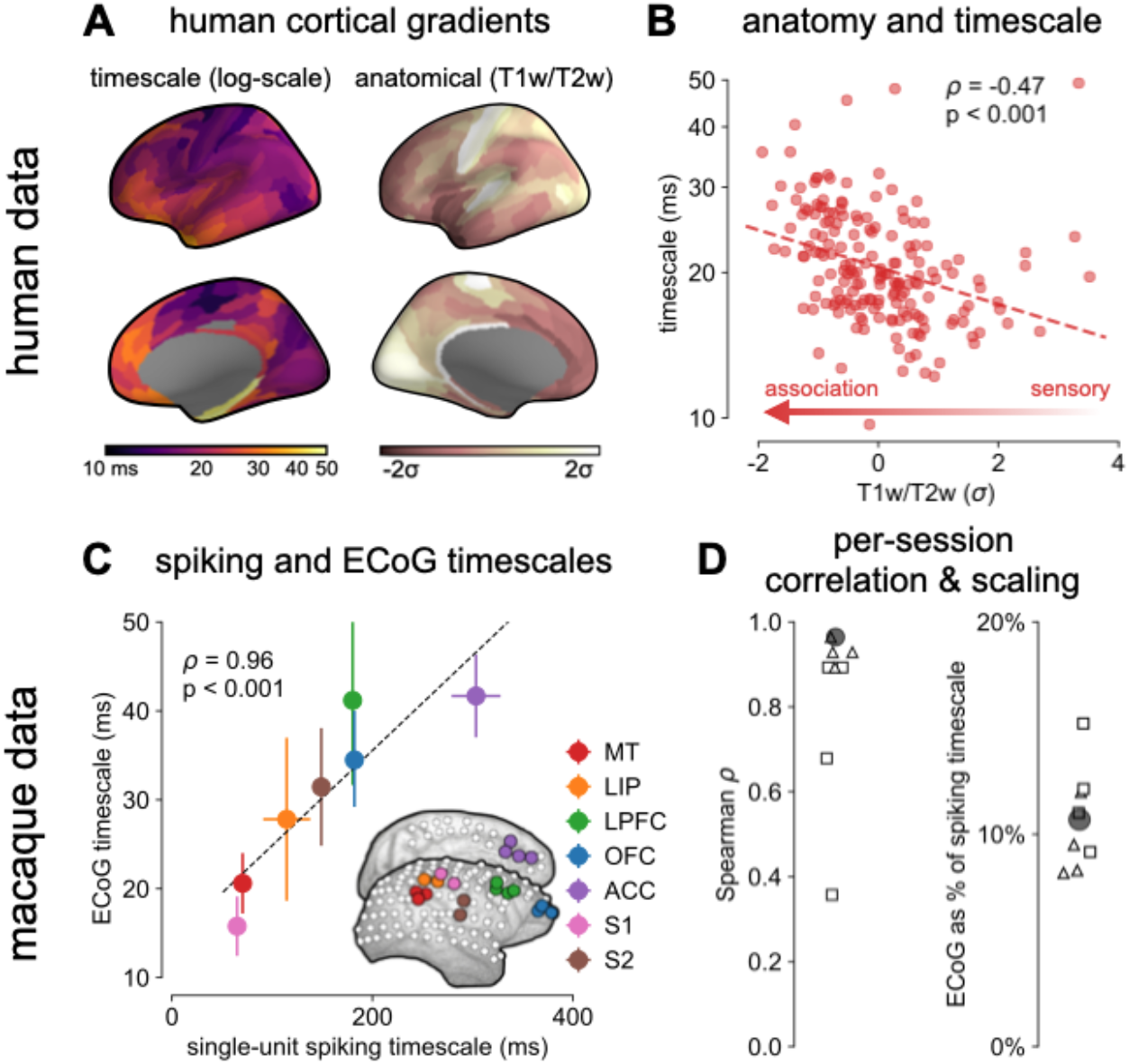
Timescale increases along the anatomical hierarchy in humans and macaques. **(A)** human cortical timescale gradient (left) falls predominantly along the rostrocaudal axis, similar to T1w/T2w ratio (right). **(B)** neuronal timescales are negatively correlated with cortical T1w/T2w, thus increasing along the anatomical hierarchy from sensory to association regions (p-value adjusted for spatial autocorrelation). **(C)** macaque ECoG timescales track published single-unit spiking timescales (Murray et al., 2014) in corresponding regions (mean±s.e.m from n=8 sessions); inset: ECoG electrode map of one animal. **(D)** ECoG-derived timescales are consistently correlated to (left), and an order of magnitude faster than (right), single-unit timescales across individual sessions. Hollow markers: individual sessions; shapes: animals; solid circles: grand average from **(C)**.

While surface ECoG recordings offer much broader spatial coverage than extracellular single-unit recordings, they are fundamentally different signals: ECoG and field potentials largely reflect integrated synaptic and other transmembrane currents across many neuronal and glial cells, rather than putative action potentials from single neurons (Buzsáki et al., 2012) (Fig. 1A, box). Considering this, we ask whether timescales measured from ECoG are related to single-unit spiking timescales along the cortical hierarchy. To test this, we extract neuronal timescales from task-free ECoG recordings in macaques and compare them to a separate dataset of single-unit spiking timescales (Murray et al., 2014) (Fig. 2C, inset; see Fig. S3 for electrode locations). Consistent with spiking timescale estimates (Murray et al., 2014; Wasmuht et al., 2018), ECoG timescales also increase along the macaque cortical hierarchy. While there is a strong correspondence between spiking and ECoG timescales (Fig. 2C; *ρ* = 0.96, *p* < 0.001)—measured from independent datasets—across the macaque cortex, ECoG-derived timescales are 10 times faster than single-unit timescales and are conserved across individual sessions (Fig. 2D). This suggests that neuronal spiking and transmembrane currents have distinct but related timescales of fluctuations, and that both are hierarchically organized along the primate cortex.

### Synaptic and ion channel genes shape timescales of neuronal dynamics

Next, we identify cellular and synaptic mechanisms underlying timescale variations across the human cortex. Theoretical accounts posit that NMDA-mediated recurrent excitation coupled with fast inhibition (Chaudhuri et al., 2015; Wang, 1999, 2008), as well as cell-intrinsic properties (Duarte and Morrison, 2019; Gjorgjieva et al., 2016; Koch et al., 1996), are crucial for shaping neuronal timescales. While *in vitro* and *in vivo* studies in model organisms (van Vugt et al., 2020; Wang et al., 2013) can test these hypotheses at the single-neuron level, causal manipulation and large-scale recording of neuronal networks embedded in the human brain is severely limited. Here, we apply an approach analogous to multimodal single-cell profiling (Bomkamp et al., 2019) and examine the transcriptomic basis of neuronal dynamics at the macroscale.

Leveraging cortex-wide bulk mRNA expression variations (Hawrylycz et al., 2012), we find that the neuronal timescale gradient overlaps with the dominant axis of gene expression across the human cortex (*ρ* = −0.60, *p* < 0.001; Fig. 3A and Fig. S4). Consistent with theoretical predictions (Fig. 3B), timescales significantly correlate with the expression of genes encoding for NMDA (GRIN2B) and GABA-A (GABRA3) receptor subunits, voltage-gated sodium (SCN1A) and potassium (KCNA3) ion channel subunits, as well as inhibitory cell-type markers (parvalbumin, PVALB), and genes previously identified to be associated with single-neuron membrane time constants (PRR5) (Bomkamp et al., 2019).

**Fig. 3.**
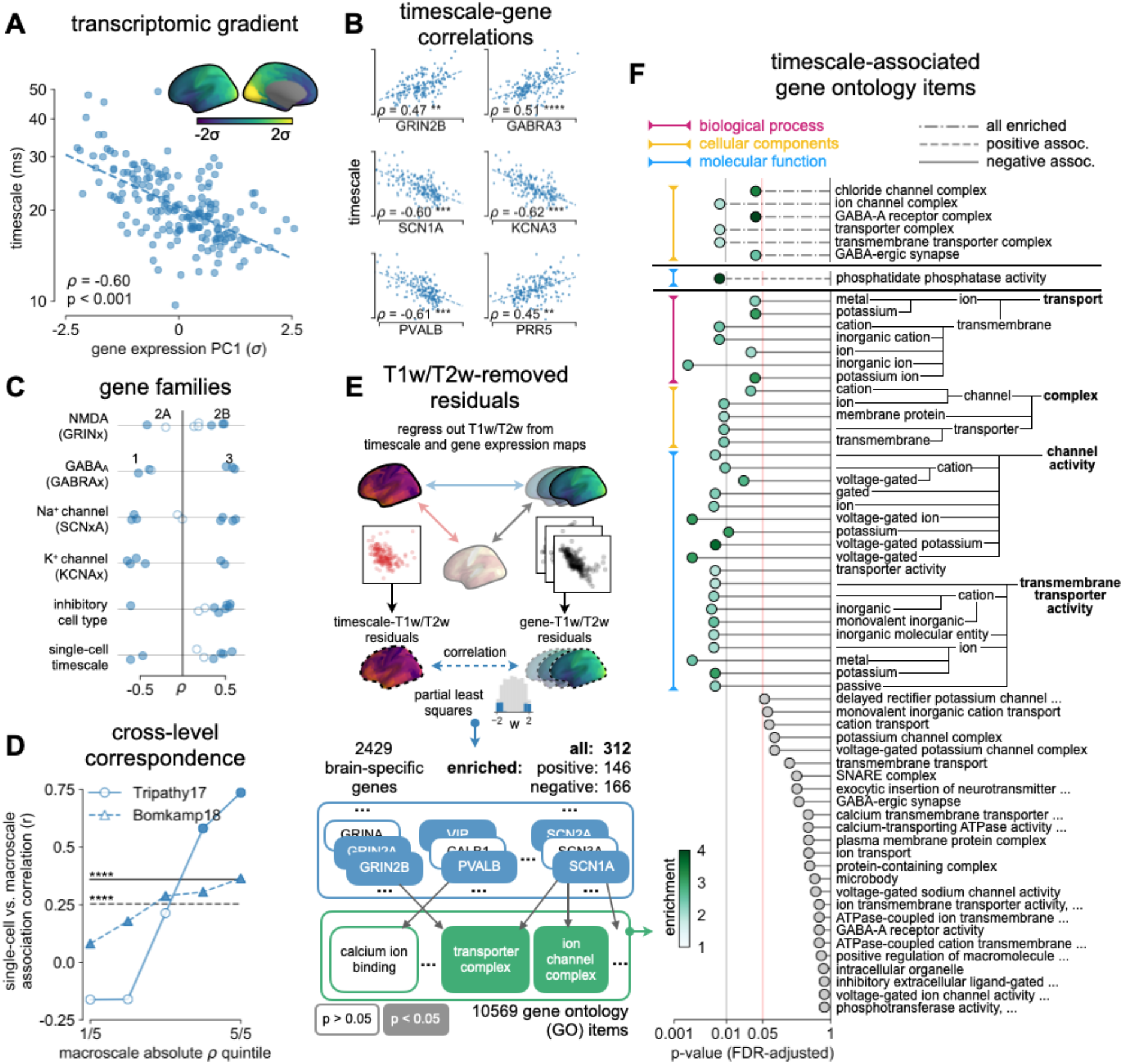
Timescale gradient is linked to expression of genes related to synaptic receptors and transmembrane ion channels across the human cortex. **(A)** timescale gradient follows the dominant axis of gene expression variation across the cortex (PC1, arbitrary direction). **(B)** timescale gradient is significantly correlated with expression of genes known to alter synaptic and neuronal membrane time constants, as well as inhibitory cell-type markers, but **(C)** members within a gene family (e.g., NMDA receptor subunits) can be both positively and negatively associated with timescales. **(D)** macroscale timescale-transcriptomic correlation captures association between RNA-sequenced expression of the same genes and single-cell timescale properties fit to patch clamp data (Bomkamp et al., 2019; Tripathy et al., 2017), and the correspondence improves for genes (separated by quintiles) that are more strongly correlated with timescale (horizontal lines: correlation across all genes from (Bomkamp et al., 2019; Tripathy et al., 2017), *ρ* = 0.36 and 0.25). **(E)** T1w/T2w gradient is regressed out from timescale and gene expression gradients, and a partial least squares (PLS) model is fit to the residual maps. Genes with significant PLS weights are submitted for gene ontology enrichment analysis. **(F)** enriched genes are primarily linked to transmembrane transporters and GABA-ergic synapses; genes specifically with strong negative associations further over-represent transmembrane ion exchange mechanisms, especially voltage-gated potassium and cation transporters. Spatial correlation p-values in **(A-C)** are adjusted for spatial autocorrelation (see Materials and Methods; asterisks in **(B,D)** indicate *p* < 0.05, 0.01, 0.005, and 0.001 respectively; filled circles in **(C,D)** indicate *p* < 0.05).

More specifically, *in vitro* electrophysiological studies have shown that, for example, increased expression of receptor subunit 2B extends the NMDA current time course (Flint et al., 1997), while 2A expression shortens it (Monyer et al., 1994). Similarly, the GABA-A receptor time constant lengthens with increasing a3:a1 subunit ratio (Eyre et al., 2012). We show that these relationships are recapitulated at the macroscale, where neuronal timescales positively correlate with GRIN2B and GABRA3 expression, and negatively correlate with GRIN2A and GABRA1. These results demonstrate that timescales of neural dynamics depend on specific receptor subunit combinations with different (de)activation timescales (Duarte et al., 2017; Gjorgjieva et al., 2016), in addition to broad excitation-inhibition interactions (Gao et al., 2017; Wang, 2002, 2020). Notably, almost all genes related to voltage-gated sodium and potassium ion channel alpha-subunits—the main functional subunits—are correlated with timescale, while all inhibitory cell-type markers except parvalbumin have strong positive associations with timescale (Fig. 3C and Fig. S5).

We further test whether single-cell timescale-transcriptomic associations are captured at the macroscale as follows: for a given gene, we can measure how strongly its expression correlates with membrane time constant parameters at the single-cell level using patch-clamp and RNA sequencing (scRNASeq) data (Bomkamp et al., 2019; Tripathy et al., 2017). Analogously, we can measure its macroscopic transcriptomic-timescale correlation using the cortical gradients above. Comparing across these two levels for all previously-identified timescale-related genes (Bomkamp et al., 2019; Tripathy et al., 2017), we find a significant correlation between the strength of association at the single-cell and macroscale levels (Fig. 3D, horizontal lines; *ρ* = 0.36 and 0.25, *p* < 0.001). Furthermore, genes with stronger associations to timescale tend to conserve this relationship across single-cell and macroscale levels (Fig. 3D, separated by macroscale correlation magnitude). Thus, the association between cellular variations in gene expression and cell-intrinsic temporal dynamics is captured at the macroscale, even though scRNAseq and microarray data represent entirely different measurements of gene expression.

While we have shown associations between cortical timescales and genes suspected to influence neuronal dynamics, these data present an opportunity to discover additional novel genes that are functionally related to timescales through a data-driven approach. However, since transcriptomic variation and anatomical hierarchy overlap along a shared macroscopic gradient (Burt et al., 2018; Huntenburg et al., 2018; Margulies et al., 2016), we cannot specify the role certain genes play based on their level of association with timescale alone: gene expression differences across the cortex first result in cell-type and connectivity differences, sculpting the hierarchical organization of cortical anatomy. Consequently, anatomy and cell-intrinsic properties jointly shape neuronal dynamics through connectivity differences (Chaudhuri et al., 2015; Demirtaş et al., 2019) and expression of ion transport proteins with variable activation timescales, respectively. Therefore, we ask whether variation in gene expression still accounts for variation in timescale beyond the principal structural gradient, and if associated genes have known functional roles in biological processes (schematic in Fig. 4E). To do this, we first remove the contribution of anatomical hierarchy by regressing out the T1w/T2w gradient from both timescale and individual gene expression gradients. We then fit partial least squares (PLS) models to simultaneously estimate regression weights for all genes (Whitaker et al., 2016), submitting those with significant associations for gene ontology enrichment analysis (GOEA) (Klopfenstein et al., 2018).

**Fig. 4.**
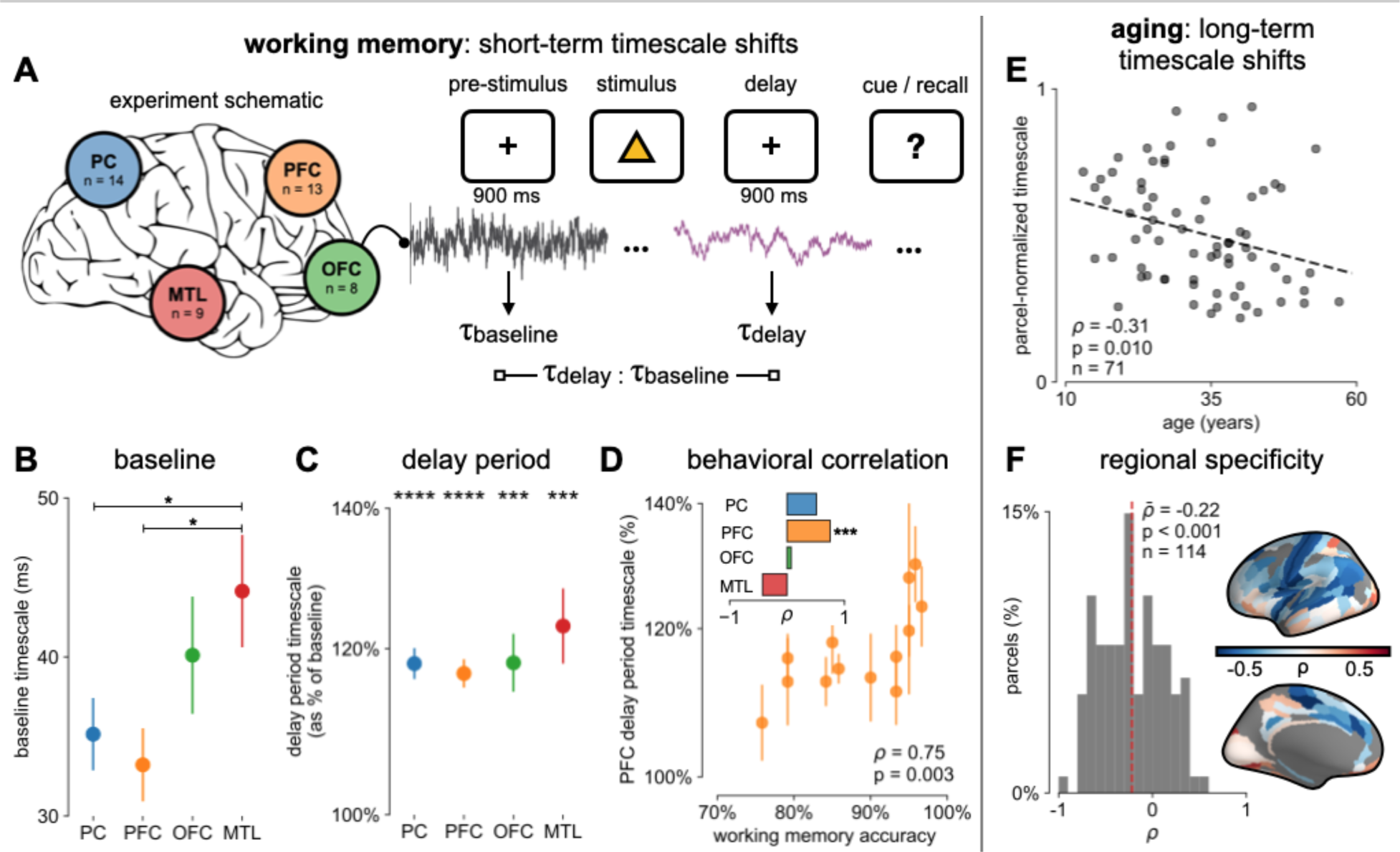
Timescales expand during working memory maintenance while tracking performance, and task-free average timescales compress in older adults. **(A)** 14 participants with overlapping intracranial coverage performed a visuospatial working memory task, with baseline (pre-stimulus) and delay period data analyzed (PC: parietal, PFC: prefrontal, OFC: orbitofrontal, MTL: medial temporal; n denotes number of subjects). **(B)** baseline timescales follow hierarchical organization within association regions (*: *p* < 0.05; mean±s.e.m. across participants). **(C)** all regions show significant timescale increase during delay period compared to baseline (***: *p* < 0.005, ****: *p* < 0.001, one-sample t-test). **(D)** PFC timescale expansion during delay periods predicts working memory accuracy across participants (mean±s.e.m. across PFC electrodes); inset: correlation between working memory accuracy and timescale change across regions. **(E)** in the MNI-iEEG dataset, participant-average cortical timescales decrease (become faster) with age. **(F)** most cortical parcels show a negative relationship between timescales and age, with the exception of parts of the visual cortex and the temporal poles (one-sample t-test, *t* = −7.04).

We find that genes highly associated with neuronal timescales are preferentially related to transmembrane ion transporter complexes, as well as GABAergic synapses and chloride channels (Fig. 4F, Table S3 and S4). When restricted to positively-associated genes only (expression increases with timescales), one functional group related to phosphatidate phosphatase activity is uncovered, including the gene PLPPR1, which has been linked to neuronal plasticity(Savaskan et al., 2004). Conversely, genes that are negatively associated with timescale are related to numerous groups involved in the construction and functioning of transmembrane transporters and voltage-gated ion channels, especially potassium and other inorganic cation transporters. The discovery of these gene ontology items suggests that inhibition (Telenczuk et al., 2017)—mediated by GABA and chloride channels—and voltage-gated potassium channels have prominent roles in shaping neuronal timescale dynamics at the macroscale level, beyond what’s expected based on the anatomical hierarchy alone.

### Timescales lengthen in working memory and shorten in aging

Finally, we investigate whether timescales are functionally dynamic and relevant for human cognition. While previous studies have shown hierarchical segregation of task-relevant information corresponding to intrinsic timescales of different cortical regions (Baldassano et al., 2017; Chien and Honey, 2020; Honey et al., 2012; Runyan et al., 2017; Sarafyazd and Jazayeri, 2019; Wasmuht et al., 2018), as well as optimal adaptation of behavioral timescales to match the environment (Ganupuru et al., 2019; Ossmy et al., 2013), evidence for functionally relevant changes in regional neuronal timescales is lacking. Here, we examine whether timescales undergo short- and long-term shifts during working memory maintenance and aging, respectively.

We first analyze human ECoG recordings where participants performed a visuospatial working memory task that requires a delayed cued response (Fig. 4A) (Johnson et al., 2018a). Neuronal timescales were extracted for pre-stimulus baseline and memory maintenance delay periods (900 ms, both stimulus-free). Replicating our previous result, we observe that baseline neuronal timescales follow a hierarchical progression across association regions (Fig. 4B). If neuronal timescales track the temporal persistence of information in a functional manner, then they should expand during delay periods. Consistent with our prediction, timescales in all regions are ~20% longer during delay periods (Fig. 4C; *p* < 0.005 for all regions). Moreover, timescale changes in the PFC are significantly correlated with behavior across participants, where longer delay-period timescales relative to baseline are associated with better working memory performance (Fig. 4D, *ρ* = 0.75, *p* = 0.003). No other spectral features in the recorded brain regions experience consistent changes from baseline to delay periods while also significantly correlating with individual performance, including the 1/f-like spectral exponent, narrowband theta (3-8 Hz), and high-frequency (high gamma; 70-100 Hz) activity power (Fig. S6).

In the long-term, aging is associated with a broad range of functional and structural changes, such as working memory impairments (Voytek et al., 2015b; Wang et al., 2011), as well as changes in neuronal dynamics (Voytek and Knight, 2015; Voytek et al., 2015b; Wang et al., 2011) and cortical structure (Pegasiou et al., 2020; de Villers-Sidani et al., 2010), such as the loss of slow-deactivating NMDA receptor subunits (Pegasiou et al., 2020). Since neuronal timescales support working memory maintenance, we specifically predict that timescales shorten across the lifespan, in agreement with the observed cognitive and structural deteriorations. To this end, we leverage the wide age range in the MNI-iEEG dataset (13-62 years old) and probe cortical timescales for each participant as a function of age. We observe that older adults have faster neuronal timescales (*ρ* = −0.31, *p* = 0.010; Fig. 4E and Fig. S7; see Materials and Methods), and that timescales shorten with age in most areas across the cortex (*t* = −7.04, *p* < 0.001). This timescale compression is especially prominent in sensorimotor, temporal, and medial frontal regions (Fig. 4F and Fig. S7). These results support our hypothesis that neuronal timescales, estimated from transmembrane current fluctuations, can rapidly shift in a functionally relevant manner, as well as slowly—over decades—in healthy aging.

## Discussion

Theoretical accounts and converging empirical evidence predict a graded variation of neuronal timescales across the human cortex (Chaudhuri et al., 2015; Huntenburg et al., 2018; Wang, 2020), which reflects functional specialization and implements hierarchical temporal processing crucial for complex cognition (Kiebel et al., 2008). This timescale gradient is thought to emerge as a consequence of cortical variations in cytoarchitecture and microcircuit connectivity, thus linking brain structure to function. In this work, we infer the timescale of non-rhythmic transmembrane current fluctuations from invasive human intracranial recordings and test those predictions explicitly.

We find that neuronal timescales vary continuously across the human cortex and coincide with the anatomical hierarchy, with timescales increasing from primary sensory and motor to association regions. Timescales inferred from macaque ECoG scale with single-unit spiking timescales, corroborating the fact that field potential signals mainly reflect fast transmembrane and synaptic currents (Buzsáki et al., 2012), whose timescales are related to, but distinct from, single-unit timescales measured in previous studies (Murray et al., 2014; Ogawa and Komatsu, 2010; Wasmuht et al., 2018). Because field potential fluctuations are driven by currents from both locally generated and distal inputs, our results raise questions on how and when these timescales interact to shape downstream spiking dynamics.

Furthermore, transcriptomic analysis demonstrates the specific roles that transmembrane ion transporters and synaptic receptors play in establishing the cortical gradient of neuronal timescales, over and above the degree predicted by the principal structural gradient alone. The expression of voltage-gated potassium channel, chloride channel, and GABAergic receptor genes, in particular, are strongly associated with the spatial variation of neuronal timescale. Remarkably, we find that electrophysiology/transcriptomic relationships discovered at the single-cell level, through patch-clamp recordings and single-cell RNA sequencing, are recapitulated at the macroscale between bulk gene expression and timescales inferred from ECoG. Our findings motivate further studies for investigating the precise roles voltage-gated ion channels and synaptic inhibition play in shaping functional neuronal timescales through causal manipulations, complementary to existing lines of research focusing on NMDA activation and recurrent circuit motifs.

Finally, we show that neuronal timescales are not static, but can change both in the short- and long-term. Transmembrane current timescales across multiple association regions, including parietal, frontal, and medial temporal cortices, increase during the delay period of a working memory task, consistent with the emergence of persistent spiking during working memory delay. Working memory performance across individuals, however, is predicted by the extent of timescale increase in the PFC only. This further suggests that behavior-relevant neural activity may be localized despite widespread task-related modulation (Pinto et al., 2019), even at the level of neuronal membrane fluctuations. In the long-term, we find that neuronal timescale shortens with age in most cortical regions, linking age-related synaptic, cellular, and connectivity changes—particularly those that influence neuronal integration timescale—to the compensatory posterior-to-anterior shift of functional specialization in healthy aging (Davis et al., 2008).

Overall, we identify consistent and converging patterns between transcriptomics, anatomy, dynamics, and function across multiple datasets of different modalities from different individuals and multiple species. As a result, evidence for these relationships can be supplemented by more targeted approaches such as imaging of receptor metabolism. Furthermore, the introduction and validation of a novel method for inferring timescales from macroscale electrophysiological recordings potentially allows for the non-invasive estimation of neuronal timescales, using widely accessible tools such as EEG and MEG (Demirtaş et al., 2019). These results open up many avenues of research for discovering potential relationships between microscale gene expression and anatomy with the dynamics of neuronal population activity at the macroscale in humans.

## Funding

Natural Sciences and Engineering Research Council of Canada (NSERC PGS-D) and Katzin Prize (to R.G.). Alexander Von Humboldt Foundation fellowship for post-doctoral researchers (to R.L.v.d.B). Deutsche Forschungsgemeinschaft (DFG) SFB936/A7/Z3 (to Tobias H. Donner). Sloan Research Fellowship (FG-2015-66057), the Whitehall Foundation (2017-12-73), the National Science Foundation under grant BCS-1736028, the NIH National Institute of General Medical Sciences grant R01GM134363-01, a UC San Diego, Shiley-Marcos Alzheimer’s Disease Research Center (ADRC): Research Training in Alzheimer’s Disease grant, and a Halιcιoğlu Data Science Institute Fellowship (to B.V.).

## Author contributions

R.G. and B.V. conception and electrophysiological data analysis. R.G. simulations, anatomical, and gene expression analysis. R.v.d.B and T.P. data preprocessing and conception of anatomical and gene expression analysis. R.G. wrote the manuscript with B.V. All authors edited the manuscript.

## Competing interests

the authors declare no competing interests.

## Data and materials availability

all data analyzed in this manuscript are from open data sources. All code used for all analyses and plots are publicly available on GitHub at https://github.com/rdgao/field-echos and https://github.com/rudyvdbrink/surface_projection. See Table S1 and S2.

## Methods

### Inferring timescale from autocorrelation and power spectral density

Consistent with previous studies, we define “neuronal timescale” as the exponential decay time constant (*τ*) of the empirical autocorrelation function (ACF), or lagged correlation (Honey et al., 2012; Murray et al., 2014). *τ* can be naively estimated to be the time it takes for the ACF to decrease by a factor of *e* when there are no additional long-term, scale-free, or oscillatory processes, or by fitting a function of the form 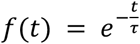 and extracting the parameter *τ*. Equivalently, the power spectral density (PSD) is the Fourier Transform of the ACF via Wiener-Khinchin theorem, and follows a Lorentzian function of the form 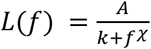 for approximately exponential-decay processes, with *χ* = 2 exactly when the ACF is solely composed of an exponential decay term, though it is often variable and in the range between 2-6 for neural time series (Haller et al., 2018; Miller et al., 2009; Podvalny et al., 2015; Voytek et al., 2015b). Timescale can be computed from the parameter *k* as 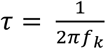, where 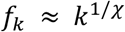 is approximated to be the frequency at which a bend or knee in the power spectrum occurs and equality holds when *χ* = 2.

### Computing power spectral density (PSD)

PSDs are estimated using a modified Welch’s method, where short-time windowed Fourier transforms (STFT) are computed from the time series, but the median is taken across time instead of the mean (in conventional Welch’s method) to minimize the effect of high-amplitude transients and artifacts (Izhikevich et al., 2018). Custom functions for this can be found in NeuroDSP (Cole et al., 2019), a published and open-source digital signal processing toolbox for neural time series (neurodsp.spectral.compute_spectrum). For simulated data, Neurotycho macaque ECoG, and MNI-iEEG datasets, we use 1-second long Hamming windows with 0.5-s overlap. To estimate single-trial PSDs for the working memory ECoG dataset (Johnson-ECoG (Johnson et al., 2018a, 2018b)), we simply apply Hamming window to 900-ms long epoched time series and compute the squared magnitude of the windowed-Fourier transform.

### Spectral parametrization – Fitting Oscillations and 1/f (FOOOF)

We apply spectral parameterization (Haller et al., 2018) to extract timescales from PSDs. Briefly, we decompose log-power spectra into a summation of narrowband periodic components—modeled as Gaussians—and an aperiodic component—modeled as a generalized Lorentzian function centered at 0 Hz (*L*(*f*) above). For inferring decay timescale, this formalism can be practically advantageous when a strong oscillatory or variable power-law (*χ*) component is present, as is often the case for neural signals. While oscillatory and power-law components can corrupt naive measurements of *τ* as time for the ACF to reach 1/*e*, they can be easily accounted for and ignored in the frequency domain as narrowband peaks and 1/f-exponent fit. We discard the periodic components and infer timescale from the aperiodic component of the PSD. For a complete mathematical description of the model, see (Haller et al., 2018).

### Simulation and validation

We simulate the aperiodic background component of neural field potential recordings as autocorrelated stochastic processes by convolving Poisson population spikes with exponentially-decaying synaptic kernels with predefined decay time constants (neurodsp.sim.sim_synaptic_current). PSDs of the simulated data are computed and parameterized as described above, and we compare the fitted timescales with their ground-truth values.

### Macaque ECoG and single unit timescales data

Macaque single-unit timescales are taken directly from values reported in Fig. 1c of (Murray et al., 2014). Whole-brain surface ECoG data (1000Hz sampling rate) is taken from the Neurotycho repository(Nagasaka et al., 2011; Yanagawa et al., 2013), with 8 sessions of 128-channel recordings from two animals (George and Chibi, 4 sessions each). Results reported in Fig. 2 are from ~10 minutes eyes-open resting periods to match the pre-stimulus baseline condition of single-unit experiments. Timescales for individual ECoG channels are extracted and averaged over regions corresponding to single-unit recording areas from(Murray et al., 2014) (Fig. 2C inset and Fig. S3), which are selected visually based on the overlapping cortical map and landmark sulci/gyri. Each region included between 2-4 electrodes (see Fig. S3C for selected ECoG channel indices for each region).

### Statistical analysis for macaque ECoG and spiking timescale

For each individual recording session, as well as the grand average, Spearman rank correlation was computed between spiking and ECoG timescales. Linear regression models were fit using the python package scipy (Virtanen et al., 2020) (scipy.stats.linregress) and the linear slope was used to compute the scaling coefficient between spiking and ECoG timescales.

### Variations in neuronal timescale, T1/T2 ratio, and mRNA expression across human cortex

The following sections describe procedures for generating the average cortical gradient maps for neuronal timescale, MR-derived T1w/T2w ratio, and gene expression from the respective raw datasets. All maps were projected onto the 180 left hemisphere parcels of Human Connectome Project’s Multimodal Parcellation (Glasser et al., 2016) (HCP-MMP1.0) for comparison, described in the individual sections. All spatial correlations are computed as Spearman rank correlations between maps. Procedure for computing statistical significance while accounting for spatial autocorrelation is described in detail below under the sections **spatial statistics** and **spatial autocorrelation modeling**.

### Neuronal timescale map

The MNI Open iEEG dataset consists of 1 minute of resting state data across 1772 channels from 106 epilepsy patients (13-62 years old, 58 males and 48 females), recorded using either surface strip/grid or stereoEEG electrodes, and cleaned of visible artifacts (Frauscher et al., 2018a, 2018b). Neuronal timescales were extracted from PSDs of individual channels, and projected from MNI voxel coordinates onto HCP-MMP1.0 surface parcellation as follows:

For each patient, timescale estimated from each electrode was extrapolated to the rest of the cortex in MNI coordinates using a Gaussian weighting function (confidence mask), *w*(*r*) = *e*^−(*r*^2^/*α*^2^)^, where r is the Euclidean distance between the electrode and a voxel, and α is the distance scaling constant, chosen here such that a voxel 4mm away has 50% weight (or, confidence). Timescale at each voxel is computed as a weighted spatial average of timescales from all electrodes (i) of that patient,

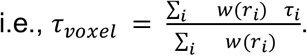

Similarly, each voxel is assigned a confidence rating that is the maximum of weights over all electrodes (*w_voxel_*(*r_min_*), of the closest electrode), i.e., a voxel right under an electrode has a confidence of 1, while a voxel 4mm away from the closest electrode has a confidence of 0.5, etc.

Timescales for each HCP-MMP parcel were then computed as the confidence-weighted arithmetic mean across all voxels that fall within the boundaries of that parcel. HCP-MMP boundary map is loaded and used for projection using NiBabel (Brett et al., 2020). This results in a 180 parcels-by-106 patients timescale matrix. A per-parcel confidence matrix of the same dimensions was computed by taking the maximum confidence over all voxels for each parcel (Fig. S1A). The average cortical timescale map (gradient) is computed by taking the confidence-weighted average at each parcel across all participants. Note that this procedure for locally thresholded and weighted average is different from projection procedures used for the mRNA and T1w/T2w data due to region-constrained and heterogeneous ECoG electrode sites across participants. While coverage is sparse and idiosyncratic in individual participants, it does not vary as a function of age, and when pooling across the entire population, 178 of 180 parcels have at least one patient with an electrode within 4mm, with the best coverage in later sensorimotor, temporal, and frontal regions (Fig. S1).

### T1w/T2w ratio map

As a measure of structural cortical hierarchy, we used the ratio between T1- and T2-weighted structural MRI, referred to as T1w/T2w map in main text, or the myelin map (Burt et al., 2018; Glasser and Van Essen, 2011). Since there is little variation in the myelin map across individuals, we used the group average myelin map of the WU-Minn HCP S1200 release (N = 1096, March 1, 2017 release) provided in HCP-MMP1.0 surface space. For correlation with other variables, we computed the median value per parcel, identical to the procedure for mRNA expression below.

### mRNA expression maps

We used the Allen Human Brain Atlas (AHBA) gene expression dataset (Hawrylycz et al., 2015, 2012) that comprised postmortem samples of 6 donors (1 female, 5 male) that underwent microarray transcriptional profiling. Spatial maps of mRNA expression were available in volumetric 2 mm isotropic MNI space, following improved nonlinear registration and whole-brain prediction using variogram modeling as implemented by (Gryglewski et al., 2018). We used whole-brain maps available from (Gryglewski et al., 2018) rather than the native sample-wise values in the AHBA database to prevent bias that could occur due to spatial inhomogeneity of the sampled locations. In total, 18114 genes were included for analyses that related to the dominant axis of expression across the genome.

We projected the volumetric mRNA expression data onto the HCP-MMP cortical surface using the HCP workbench software (v1.3.1 running on Windows OS 10) with the “enclosing” method, and custom MATLAB code (github.com/rudyvdbrink/surface_projection). The enclosing method extracts for all vertices on the surface the value from enclosing voxels in the volumetric data. Alternative projection methods such as trilinear 3D linear interpolation of surrounding voxels, or ribbon mapping that constructs a polyhedron from each vertex’s neighbors on the surface to compute a weighted mean for the respective vertices, yielded comparable values, but less complete cortical coverage. Moreover, the enclosing method ensured that no transformation of the data (non-linear or otherwise) occurred during the projection process and thus the original values in the volumetric data were preserved.

Next, for each parcel of the left hemisphere in HCP-MMP, we extracted the median vertex-wise value. We used the median rather than the mean because it reduced the contribution of outliers in expression values within parcels. Vertices that were not enclosed by voxels that contained data in volumetric space were not included in the parcel-wise median. This was the case for 539 vertices (1.81% of total vertices). Linear interpolation across empty vertices prior to computing median parcel-wise values yielded near-identical results (*r* = 0.95 for reconstructed surfaces). Lastly, expression values were mean and variance normalized across parcels to facilitate visualization. Normalization had no effect on spatial correlation between gene expression and other variables since the spatial distribution of gene expression was left unaltered.

### Spatial statistics

All correlations between spatial maps (timescale, T1w/T2w, gene principal component, and individual gene expressions) were computed using Spearman rank correlation. As noted in (Burt et al., 2018, 2020; Vos de Wael et al., 2020), neural variables vary smoothly and continuously across the cortical surface, violating the assumption of independent samples. As a result, when correlating two variables each with non-trivial spatial autocorrelation, the naive p-value is artificially lowered since it is compared against an inappropriate null hypothesis, i.e., randomly distributed or shuffled values across space. Instead, a more appropriate null hypothesis introduces spatial autocorrelation-preserving null maps, which destroys any potential correlation between two maps while respecting their spatial autocorrelations. For all spatial correlation analyses, we generated N = 1000 null maps of one variable (timescale map unless otherwise noted), and the test statistic, Spearman correlation (ρ), is computed against the other variable of interest to build the null distribution. Two-tailed significance is then computed as the proportion of the null distribution that is less extreme than the empirical correlation value. All regression lines were computed by fitting a linear regression to log-timescale and the structural feature maps.

### Spatial autocorrelation modeling

To generate spatial autocorrelation-preserving null maps, we used Moran’s Spectral Randomization (MSR) (Wagner and Dray, 2015) from the python package BrainSpace (Vos de Wael et al., 2020). Details of the algorithm can be found in the above references. Briefly, MSR performs eigendecomposition on a spatial weight matrix of choice, which is taken here to be the inverse average geodesic distance matrix between all pairs of parcels in HCP-MMP1.0. The eigenvectors of the weight matrix are then used to generate randomized null feature maps that preserves the autocorrelation of the empirical map. We used the singleton procedure for null map generation. All significance values reported (Fig. 2B, Fig. 3A-C) were adjusted using the above procedure.

We also compare two other methods of generating null maps: spatial variogram fitting (Burt et al., 2020) and spin permutation (Alexander-Bloch et al., 2018). Null maps were generated for timescale using spatial variogram fitting, while for spin permutation they were generated for vertex-wise T1w/T2w and gene PC1 maps before parcellation, so as to preserve surface locations of the parcellation itself. All methods perform similarly, producing comparable spatial autocorrelation in the null maps, assessed using spatial variogram, as well as null distribution of spatial correlation coefficients between timescale and T1w/T2w (Fig. S2).

### Principal Component Analysis (PCA) of gene expression

We used scikit-learn (Pedregosa, 2011) PCA (sklearn.decomposition.PCA) to identify the dominant axes of gene expression variation across the entire AHBA dataset, as well as for brain-specific genes. PCA was computed on the variance-normalized average gene expression maps, *X*, an N x P matrix where N = 18114 (or N = 2429 brain-specific) genes, and P = 180 cortical parcels. Briefly, PCA factorizes *X* such that *X* = *USV^T^*, where *U* and *V* are unitary matrices of dimensionality N x N and P x P, respectively. *S* is the same dimensionality as *X* and contains non-negative descending eigenvalues on its main diagonal (Λ). Columns of *V* are defined as the principal components (PCs), and the dominant axis of gene expression is then defined as the first column of V, whose proportion of variance explained in the data is the first element of Λ divided by the sum over Λ. Results for PC1 and PC2-10 are shown in Fig. 3A and Fig. S4, respectively.

### Selection of brain-specific genes

Similar to (Burt et al., 2018; Fagerberg et al., 2014; Genovese et al., 2016), N=2429 brain-specific genes were selected based on the criteria that expression in brain tissues were 4 times higher than the median expression across all tissue types, using Supplementary Dataset 1 of (Fagerberg et al., 2014). PC1 result shown in Fig. 3A is computed from brain-specific genes, though findings are identical when using all genes (*ρ* = −0.56 with timescale map, Fig. S4).

### Comparison of timescale-transcriptomic association with single-cell timescale genes

Single-cell timescale genes were selected based on data from Table S3 and Online Table 1 of (Bomkamp et al., 2019; Tripathy et al., 2017), respectively. Using single-cell RNA sequencing data and patch-clamp recordings from transgenic mice cortical neurons, these studies identified genes whose expression significantly correlated with electrophysiological features derived from generalized linear integrate and fire (GLIF) model fits. We selected genes that were significantly correlated to membrane time constant (*tau*), input resistance (*Rin* or *ri*), or capacitance (*Cm* or *cap*) in the referenced data tables, and extracted the level of association between gene expression and those electrophysiological feature (correlation ‘DiscCorr’ in (Tripathy et al., 2017) and linear coefficient “beta_gene” in (Bomkamp et al., 2019)).

To compare timescale-gene expression association at the single-cell and macroscale level, we correlated the single-cell associations extracted above with the spatial correlation coefficient (macroscale ρ) between ECoG timescale and AHBA microarray expression data for those same genes, restricting to genes with *p* < 0.05 for macroscale correlation (results identical for non-restrictive gene set). Overall association for all genes, as well as split by quintiles of their absolute macroscale correlation coefficient, are shown in Fig. 3D. Example “single-cell timescale” genes shown in Fig. 3B,C are genes showing the highest correlations with those electrophysiology features reported in Table 2 of (Bomkamp et al., 2019).

### T1w/T2w-removed timescale and gene expression residual maps

To remove anatomical hierarchy as a potential mediating variable in timescale-gene expression relationships, we linearly regress out the T1w/T2w map from the (log) timescale map and individual gene expression maps. T1w/T2w was linearly fit to log-timescale, and the error between T1w/T2w-predicted timescale and empirical timescale was extracted (residual); this identical procedure was applied to every gene expression map to retrieve the gene residuals. Spatial autocorrelation-preserving null residual maps were similarly created using MSR.

### Partial least squares regression model

Due to multicollinearity in the high-dimensional gene expression dataset (many more genes than parcels), we fit a partial least squares model to the timescale map with one output dimension (sklearn.cross_decomposition.PLSRegression) to estimate regression coefficient for all genes simultaneously, resulting in N=18114 (or N=2429 brain-specific) PLS weights(Vértes et al., 2016; Whitaker et al., 2016). To determine significantly associated (or, “enriched”) genes, we repeated the above PLS-fitting procedure 1000 times but replaced the empirical timescale map (or residual map) with null timescale maps (or residual maps) that preserved its spatial autocorrelation. Genes whose absolute empirical PLS weight that was greater than 95% of its null weight distribution was deemed to be enriched, and submitted for gene ontology enrichment analysis.

### Gene ontology enrichment analysis (GOEA)

The Gene Ontology (GO) captures hierarchically structured relationships between GO items representing aspects of biological processes (BP), cellular components (CC), or molecular functions (MF). For example, “synaptic signaling”, “chemical synaptic transmission”, and “glutamatergic synaptic transmission” are GO items with increasing specificity, with smaller subsets of genes associated with each function. Each GO item is annotated with a list of genes that have been linked to that particular process or function. GOEA examines the list of enriched genes from above to identify GO items that are more associated with those genes than expected by chance. We used GOATOOLS (Klopfenstein et al., 2018) to perform GOEA programmatically in python.

The list of unranked genes with significant empirical PLS weights was submitted for GOEA as the “study set”, while either the full ABHA list or brain-specific gene list was used as the “reference set”. The output of GOEA is a list of GO terms with annotated genes that are enriched or purified (i.e., preferentially appearing or missing in the study list, respectively) more often than by chance, determined by Fisher’s exact test.

Enrichment ratio is defined as follows: given a reference set with *N* total genes, and *n* were found to be significantly associated with timescale (in the study set), for a single GO item with *B* total genes annotated to it, where b of them overlap with the study set, then 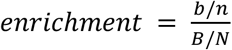. Statistical significance is adjusted for multiple comparisons following Benjamini-Hochberg procedure (false discovery rate q-value reported in Fig. 3F), and all significant GO items (q < 0.05) are reported in Fig. 3F, in addition to some example items that did not pass significance threshold. For a detailed exposition, see (Bauer, 2017). Fig. 3F shows results using brain-specific genes. The GO items that are significantly associated are similar when using the full gene set, but typically with larger q-values (Tables S3 and S4) due to a mucher larger set of (non-brain-specific) genes.

### Working memory ECoG data and analysis

The CRCNS fcx-2 and fcx-3 datasets include 17 intracranial ECoG recordings in total from epilepsy patients (10 and 7, respectively) performing the same visuospatial working memory task (Johnson, 2018, 2019; Johnson et al., 2018a, 2018b). Subject 3 (s3) from fcx-2 was discarded due to poor data quality upon examination of trial-averaged PSDs (high noise floor near 20 Hz), while s5 and s7 from fcx-3 correspond to s5 and s8 in fcx-2 and were thus combined. Together, data from 14 unique participants (22-50 years old, 5 female) were analyzed, with variable and overlapping coverage in parietal cortex (PC, n=14), prefrontal cortex (PFC, n=13), orbitofrontal cortex (OFC, n=8), and medial temporal lobe (MTL, n=9). Each channel was annotated as belonging to one of the above macro regions.

Experimental setup is described in (Johnson, 2018, 2019; Johnson et al., 2018a, 2018b) in detail. Briefly, following a 1-second pre-trial fixation period (baseline), subjects were instructed to focus on one of two stimulus contexts (“identity” or “relation” information). Then two shapes were presented in sequence for 200 ms each. After a 900 or 1150 ms jittered precue delay (delay1), the test cue appeared for 800 ms, followed by another post-cue delay period of the same length (delay2). Finally, the response period required participants to perform a 2-alternative forced choice test based on the test cue, which varied based on trial condition. For our analysis, we collapsed across the stimulus context conditions and compared neuronal timescales during the last 900 ms of baseline and delay periods from the epoched data, which were free of visual stimuli, in order to avoid stimulus-related event-related potential effects. Behavioral accuracy for each experimental condition was reported for each participant, and we average across both stimulus context conditions to produce a single working memory accuracy per participant.

Single-trial power spectra were computed for each channel as the squared magnitude of the Hamming-windowed Fourier Transform. We used 900 ms of data in all 3 periods (pre-trial, delay1, and delay2). Timescales were estimated by applying spectral parameterization as above, and the two delay-period estimates were averaged to produce a single delay period value. For comparison, we computed single-trial theta (3-8 Hz) and high-frequency activity (high gamma (Mukamel et al., 2005), 70-100 Hz) powers as the mean log-power within those frequency bins, as well as spectral exponent (χ). Single-trial timescale difference between delay and baseline was calculated as the difference of the log timescales due to the non-normal distribution of single-trial timescale estimates. All other neural features were computed by subtracting baseline from the delay period.

All neural features were then averaged across channels within the same regions, then trials, for each participant, to produce per-participant region-wise estimates, and finally averaged across all participants for the regional average in Fig. 4B,C. One-sample two-sided t-tests were used to determine the statistical significance of timescale change in each region (Fig. 4C), where the null hypothesis was no change between baseline and delay periods (i.e., delay is 100% of baseline). Spearman rank correlation was used to determine the relationship between neural activity (timescale; theta; high-frequency; χ) change and working memory accuracy across participants (Fig. 4D, Fig. S6).

### Per-subject average cortical timescale across age

Since electrode coverage in the MNI-iEEG dataset is sparse and non-uniform across participants (Fig. S1), simply averaging across parcels within individuals to estimate an average cortical timescale per participant confounds the effect of age with the spatial effect of cortical hierarchy. Therefore, we instead first normalize each parcel by its max value across all participants before averaging within participants, excluding those with fewer than 10 valid parcels (71 of 106 subjects remaining; results hold for a range of threshold values; Fig. S7B). Spearman rank correlation was used to compute the association between age and average cortical timescale.

### Age-timescale association for individual parcels

Each cortical parcel had a variable number of participants with valid timescale estimates above the consistency threshold, so we compute Spearman correlation between age and timescale for each parcel, but including only those with at least 5 participants (114 of 180 parcels, result holds for a range of threshold values; Fig. S7C). Spatial effect of age-timescale variation is plotted in Fig. 4F, where parcels that did not meet the threshold criteria are greyed out. Mean age-timescale correlation from individual parcels was significantly negative under one-sample t-test.

## Supplemental Information

**Fig. S1.**
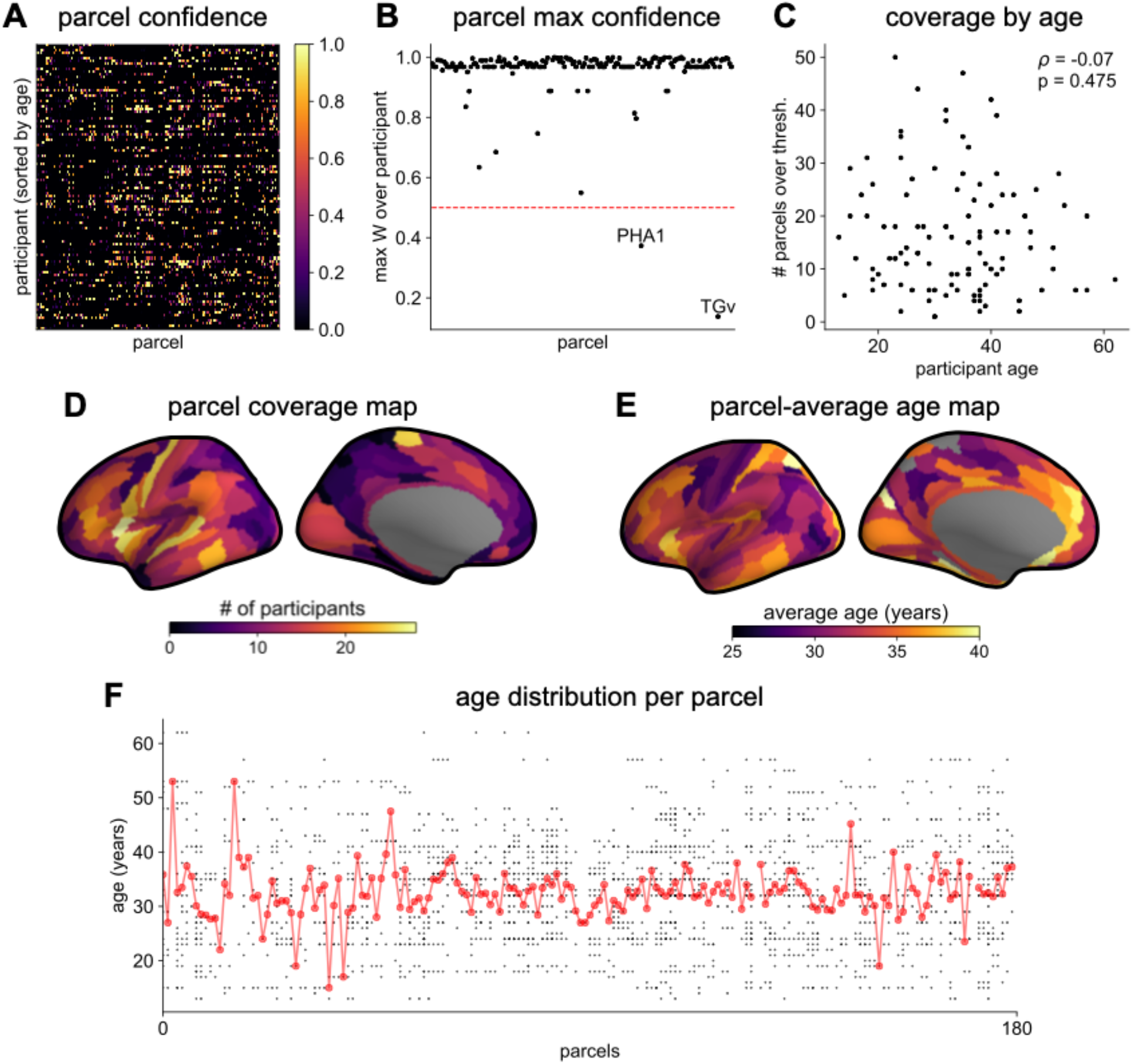
MNI-iEEG dataset coverage. **(A)**, per-parcel Gaussian-weighted mask values showing how close the nearest electrode was to a given HCP-MMP1.0 parcel for each participant. Brighter means closer, 0.5 corresponds to the nearest electrode being 4 mm away. **(B)** maximum weight for each parcel across all participants. Most parcels have electrodes very close by across the entire participant pool. **(C)** the number of HCP-MMP parcels each participant has above the confidence threshold of 0.5 is uncorrelated with age. **(D)** number of participants with confidence above threshold at each parcel. Sensorimotor, frontal, and lateral temporal regions have the highest coverage. **(E)** average age of participants with confidence above threshold at each parcel. **(F)** age distribution of participants with confidence above threshold at each parcel. Average age per parcel (red line) is relatively stable while age distribution varies from parcel to parcel.

**Fig. S2.**
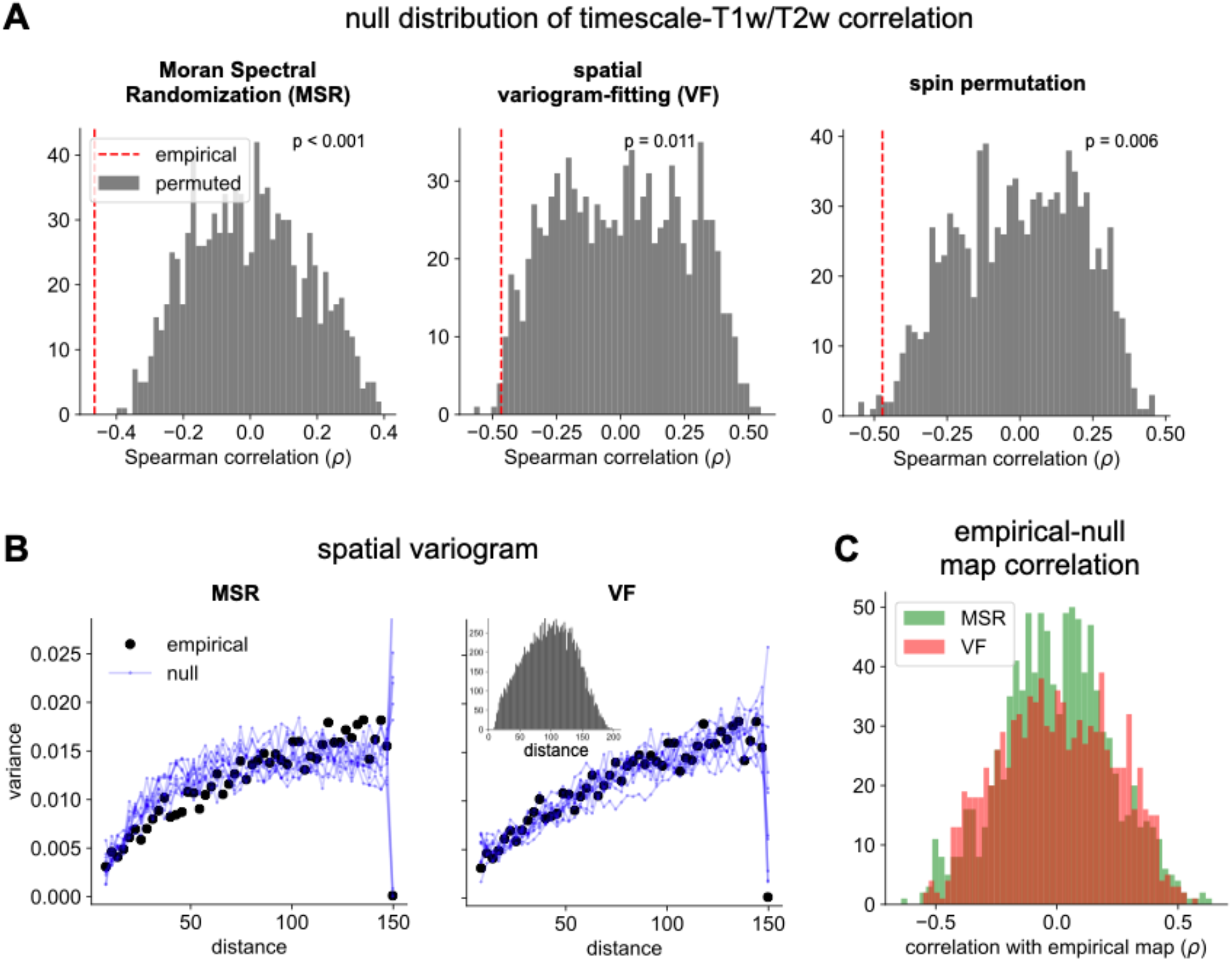
Comparison of spatial autocorrelation-preserving null map generation methods. **(A)** distributions of Spearman correlation values between empirical T1w/T2w map and 1000 spatial-autocorrelation preserving null timescale maps generated using Moran Spectral Randomization (MSR), spatial variogram fitting (VF), and spin permutation. Red dashed line denotes correlation between empirical timescale and T1w/T2w maps, p-values indicate two-tailed significance, i.e., proportion of distribution with values more extreme than empirical correlation. **(B)** spatial variogram for empirical timescale map (black) and 10 null maps (blue) generated using MSR and VF. Inset shows distribution of distances between pairs of HCP-MMP parcels. **(C)** distribution of Spearman correlations between empirical and 1000 null timescale maps generated using MSR (green) and VF (red), showing similar levels of correlation between empirical and null maps for both methods.

**Fig. S3.**
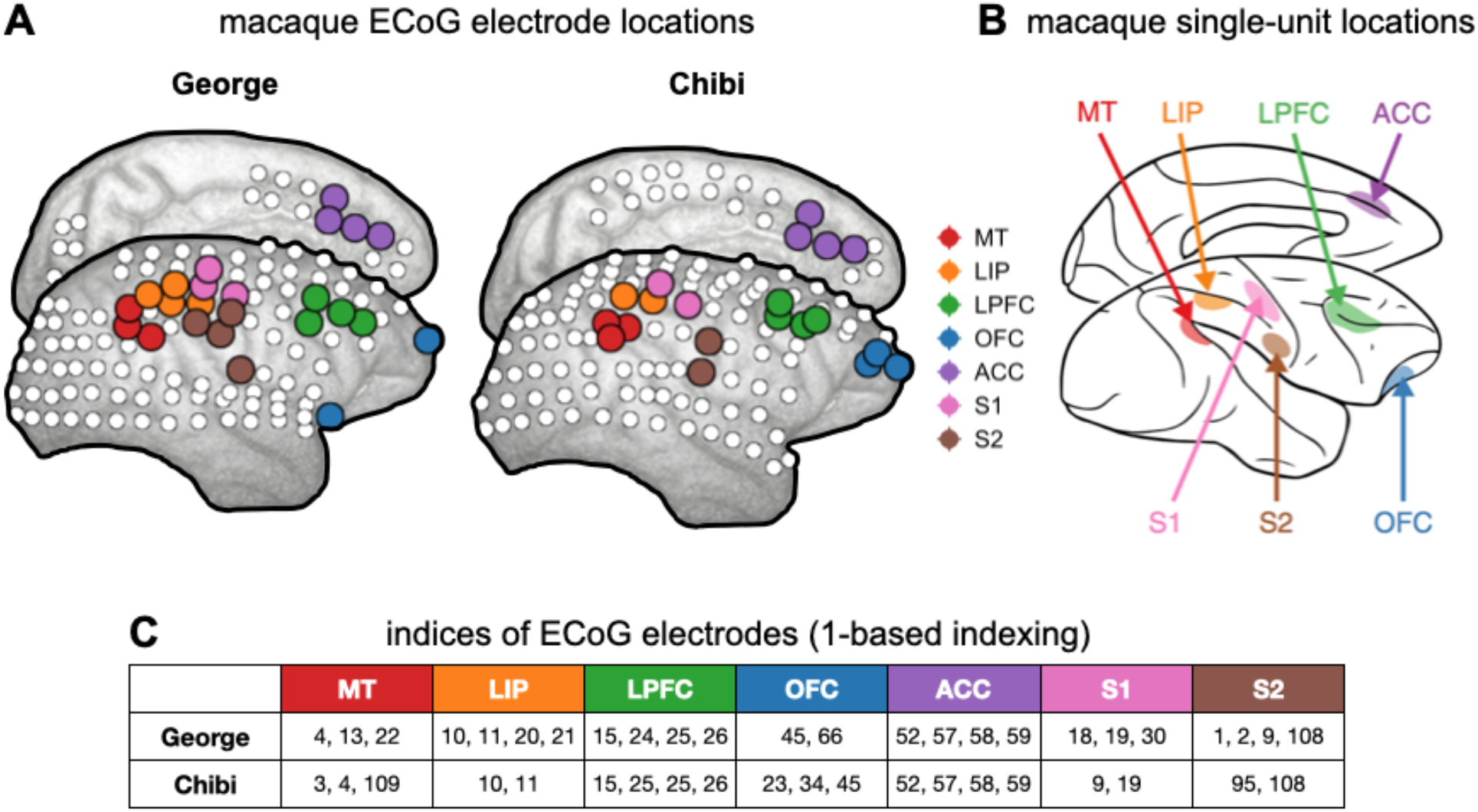
Macaque ECoG and single-unit coverage. **(A)** locations of 180-electrode ECoG grid from 2 animals in the Neurotycho dataset, colors correspond to locations used for comparison with single-unit timescales. **(B)** single-unit recording locations from Fig. 1a of (Murray et al., 2014). **(C)** electrode indices of the sampled areas from the two animals, corresponding to those colored in **(A)**.

**Fig. S4.**
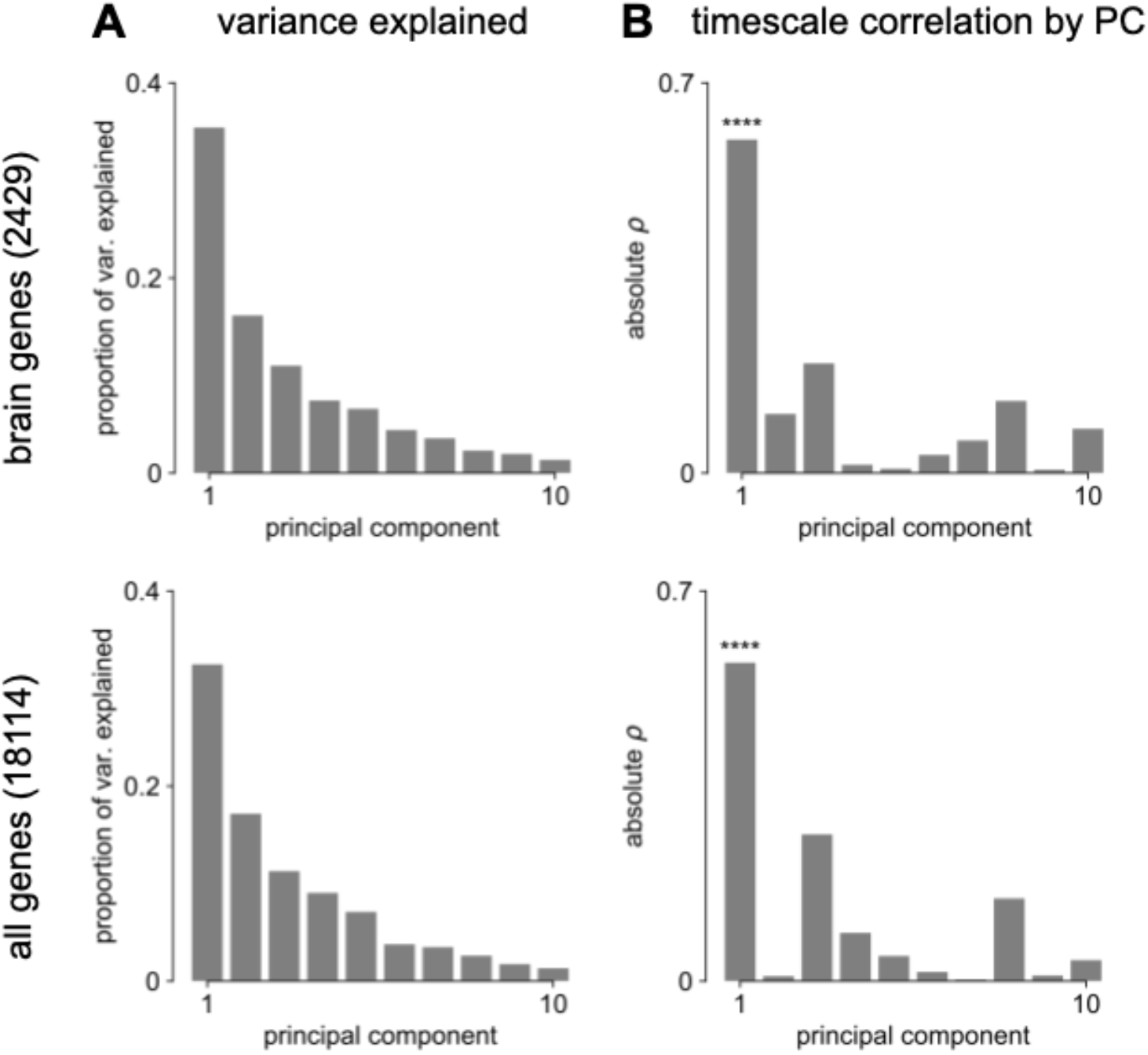
Transcriptomic PCA results. **(A)** proportion of variance explained by the top 10 principal components (PCs) of brain-specific genes (top) and all AHBA genes (bottom). **(B)** absolute Spearman correlation between timescale map and top 10 PCs from brain-specific or full gene dataset. Asterisks indicate resampled significance while accounting for spatial autocorrelation, **** indicate *p* < 0.001.

**Fig. S5.**
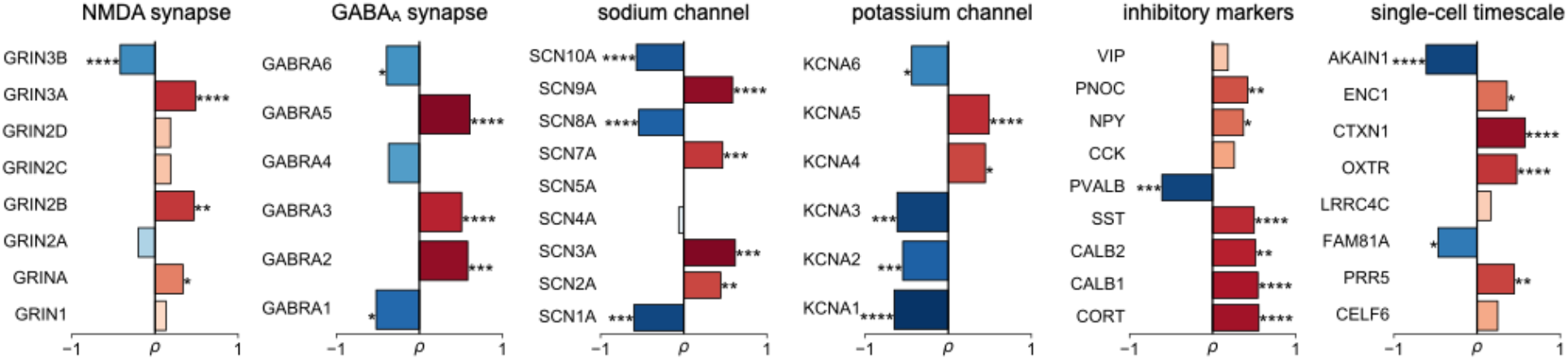
Individual gene correlations from Fig. 3C with gene symbols labeled, and grouped into functional families.

**Fig. S6.**
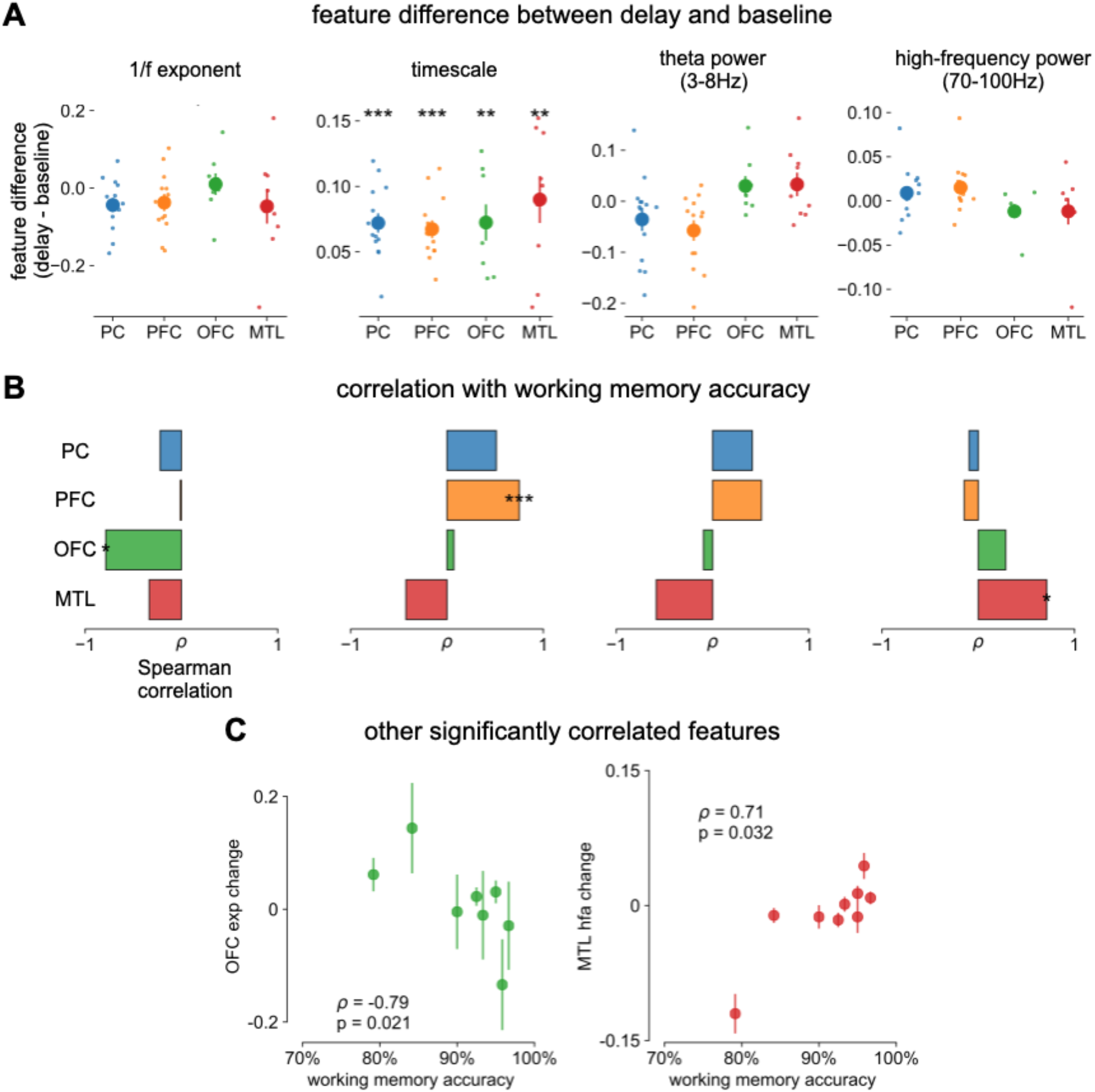
Spectral correlates of working memory performance. **(A)** difference between delay and baseline periods for 1/f-exponent, timescale (same as main Fig. 4C but absolute units on y-axis, instead of percentage), theta power, and high-frequency power. **(B)** Spearman correlation between spectral feature difference and working memory accuracy across participants, same features as in **(A)**. * *p* < 0.05, ** *p* < 0.01, *** *p* < 0.005 in **(A, B)**. **(C)** scatter plot of other significantly correlated spectral features from **(B)**.

**Fig. S7.**
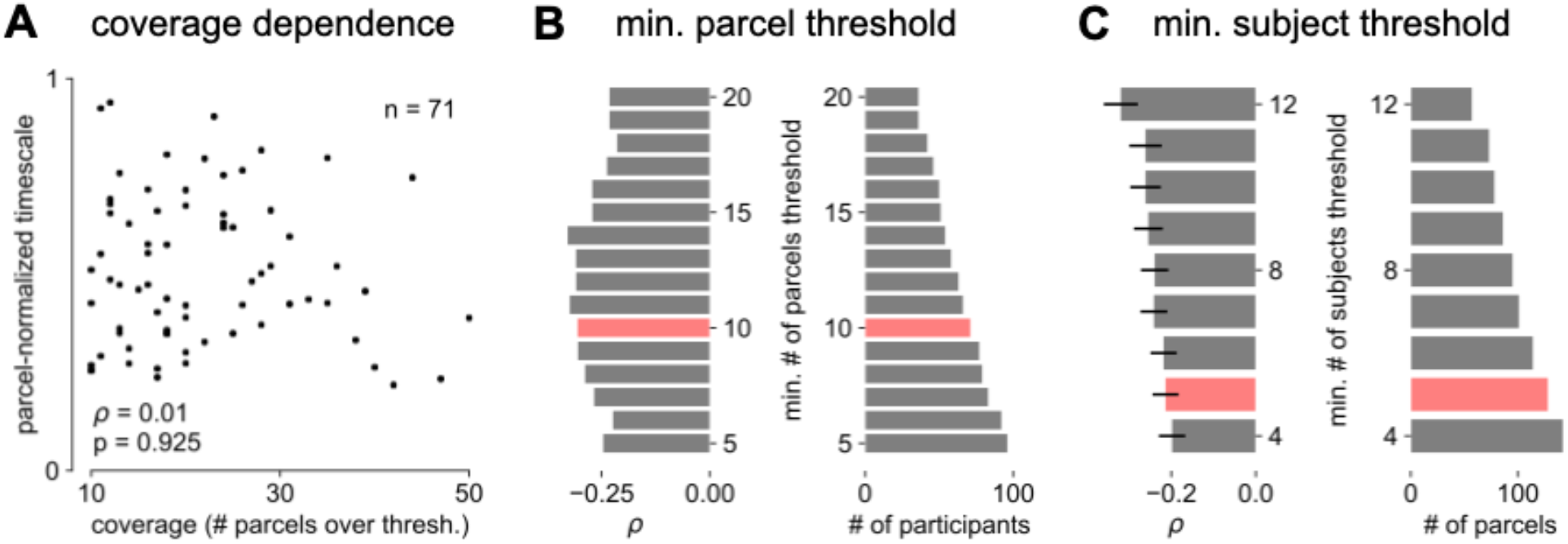
Parameter sensitivity for timescale-aging analysis. **(A)** cortex-averaged timescale is independent of parcel coverage across participants. **(B)** sensitivity analysis for the number of valid parcels a participant must have in order to be included in analysis for main Fig. 4E (red). As threshold increases (more stringent), fewer participants satisfy the criteria (right) but correlation between participant age and timescale remains robust (left). **(C)** sensitivity analysis for the number of valid participants a parcel must have in order to be included in analysis for main Fig. 4F. As threshold increases (more stringent), fewer parcels satisfy the criteria (right) but average correlation across all parcels remains robust (left, error bars denote s.e.m of distribution as in Fig. 4F).

**Table S1.** Summary of open-access datasets used

| Data | Ref. | Specific Source/<br>Format Used | Relevant Figures |
| --- | --- | --- | --- |
| MNI Open iEEG Atlas | (Frauscher et al., 2018a, 2018b) |  | Fig. 2A,B,<br>Fig. 3,<br>Fig. 4 |
| Tw1/T2w map<br>Human Connectome<br>Project | (Glasser and Van Essen, 2011; Glasser et al., 2016) | Release S1200,<br>March 1, 2017 | Fig. 2A,B,<br>Fig. 3D-F |
| Neurotycho macaque<br>ECoG | (Nagasaka et al., 2011;<br>Yanagawa et al., 2013) | Anesthesia datasets,<br>propofol and<br>ketamine (Chibi and<br>Gerge) | Fig. 2C,D |
| Macaque single-unit<br>timescales | (Murray et al., 2014) | Fig. 1 of reference | Fig. 2C,D |
| Whole-cortex interpolated<br>Allen Brain Atlas human<br>gene expression | (Gryglewski et al., 2018;<br>Hawrylycz et al., 2012) | Interpolated maps<br>downloadable from<br>http://www.meduniwien.ac.at/neuroimaging/mRNA.html | Fig. 3 |
| Single-cell timescale-related genes | (Bomkamp et al., 2019;<br>Tripathy et al., 2017) | Table S3 from<br>(Tripathy et al.,<br>2017), Online Table<br>1 from (Bomkamp et<br>al., 2019) | Fig. 3C,D |
| Human working memory<br>ECoG | (Johnson, 2018, 2019;<br>Johnson et al., 2018a,<br>2018b) | CRCNS fcx-2 and<br>fcx-3 | Fig. 4A-D |

**Table S2.** Reproducing figures from code repository

| All IPython notebooks: https://github.com/rdgao/field-echos/tree/master/notebooks |  |
| --- | --- |
| Notebook | Results |
| 2a_sim_method_schematic.ipynb | **simulations:** Fig. 1B-E, Fig. S3 |
| 2b_viz_NeuroTycho-SU.ipynb | **macaque timescales:** Fig. 2C,D |
| 3_viz_human_structural.ipynb | **human timescales vs. T1w/T2w and gene expression:** Fig. 2A,B, Fig. 3, Fig. S1, S4, S5; Table. S3 |
| 4b_viz_human_wm.ipynb | **human working memory:** Fig. 4A-D, Fig. S6 |
| 4a_viz_human_aging.ipynb | **human aging:** Fig. 4E,F, Fig. S7 |
| supp_spatialautocorr.ipynb | **spatial autocorrelation-preserving nulls:** Fig. S2 |
| Projection of T1w/T2w and gene expression maps from MNI volumetric coordinates to HCP-MMP1.0 can be found: https://github.com/rudyvdbrink/Surface_projection |  |

**Table S3.** Significant items from brain-specific GOEA (Fig. 3F)

| gene association | ID | e/p | ontology | name | enrichment ratio | p-value (FDR-adjusted) |
| --- | --- | --- | --- | --- | --- | --- |
| all | GO:0034702 | e | CC | ion channel complex | 1.959 | 0.008 |
| all | GO:1902495 | e | CC | transmembrane transporter complex | 1.91 | 0.008 |
| all | GO:1990351 | e | CC | transporter complex | 1.91 | 0.008 |
| all | GO:0098982 | e | CC | GABA-ergic synapse | 2.497 | 0.038 |
| all | GO:1902711 | e | CC | GABA-A receptor complex | 4.541 | 0.038 |
| all | GO:0034707 | e | CC | chloride channel complex | 3.385 | 0.038 |
| pos | GO:0008195 | e | MF | phosphatidate phosphatase activity | 13.864 | 0.007 |
| neg | GO:0098660 | e | BP | inorganic ion transmembrane transport | 2.515 | 0.002 |
| neg | GO:0098662 | e | BP | inorganic cation transmembrane transport | 2.529 | 0.007 |
| neg | GO:0098655 | e | BP | cation transmembrane transport | 2.439 | 0.007 |
| neg | GO:0034220 | e | BP | ion transmembrane transport | 2.057 | 0.03 |
| neg | GO:0030001 | e | BP | metal ion transport | 2.239 | 0.036 |
| neg | GO:0071805 | e | BP | potassium ion transmembrane transport | 3.122 | 0.036 |
| neg | GO:0006813 | e | BP | potassium ion transport | 3.081 | 0.037 |
| neg | GO:1902495 | e | CC | transmembrane transporter complex | 2.334 | 0.009 |
| neg | GO:1990351 | e | CC | transporter complex | 2.334 | 0.009 |
| neg | GO:0034702 | e | CC | ion channel complex | 2.36 | 0.009 |
| neg | GO:0098796 | e | CC | membrane protein complex | 2.063 | 0.009 |
| neg | GO:0034703 | e | CC | cation channel complex | 2.379 | 0.03 |
| neg | GO:0005244 | e | MF | voltage-gated ion channel activity | 3.081 | 0.002 |
| neg | GO:0022832 | e | MF | voltage-gated channel activity | 3.081 | 0.002 |
| neg | GO:0046873 | e | MF | metal ion transmembrane transporter activity | 2.453 | 0.002 |
| neg | GO:0022890 | e | MF | inorganic cation transmembrane transporter activity | 2.24 | 0.005 |
| neg | GO:0005216 | e | MF | ion channel activity | 2.289 | 0.006 |
| neg | GO:0008324 | e | MF | cation transmembrane transporter activity | 2.173 | 0.006 |
| neg | GO:0015318 | e | MF | inorganic molecular entity transmembrane transporter activity | 2.04 | 0.006 |
| neg | GO:0015077 | e | MF | monovalent inorganic cation transmembrane transporter activity | 2.535 | 0.006 |
| neg | GO:0015075 | e | MF | ion transmembrane transporter activity | 2.024 | 0.006 |
| neg | GO:0015079 | e | MF | potassium ion transmembrane transporter activity | 3.041 | 0.006 |
| neg | GO:0005215 | e | MF | transporter activity | 1.883 | 0.006 |
| neg | GO:0022857 | e | MF | transmembrane transporter activity | 1.906 | 0.006 |
| neg | GO:0022836 | e | MF | gated channel activity | 2.301 | 0.006 |
| neg | GO:0015267 | e | MF | channel activity | 2.191 | 0.006 |
| neg | GO:0022803 | e | MF | passive transmembrane transporter activity | 2.191 | 0.006 |
| neg | GO:0005249 | e | MF | voltage-gated potassium channel activity | 3.658 | 0.006 |
| neg | GO:0005261 | e | MF | cation channel activity | 2.353 | 0.009 |
| neg | GO:0005267 | e | MF | potassium channel activity | 3.058 | 0.011 |
| neg | GO:0022843 | e | MF | voltage-gated cation channel activity | 2.744 | 0.022 |

e/p: enriched or purified; BP: biological process; CC: cellular components; MF: molecular function

**Table S4.**
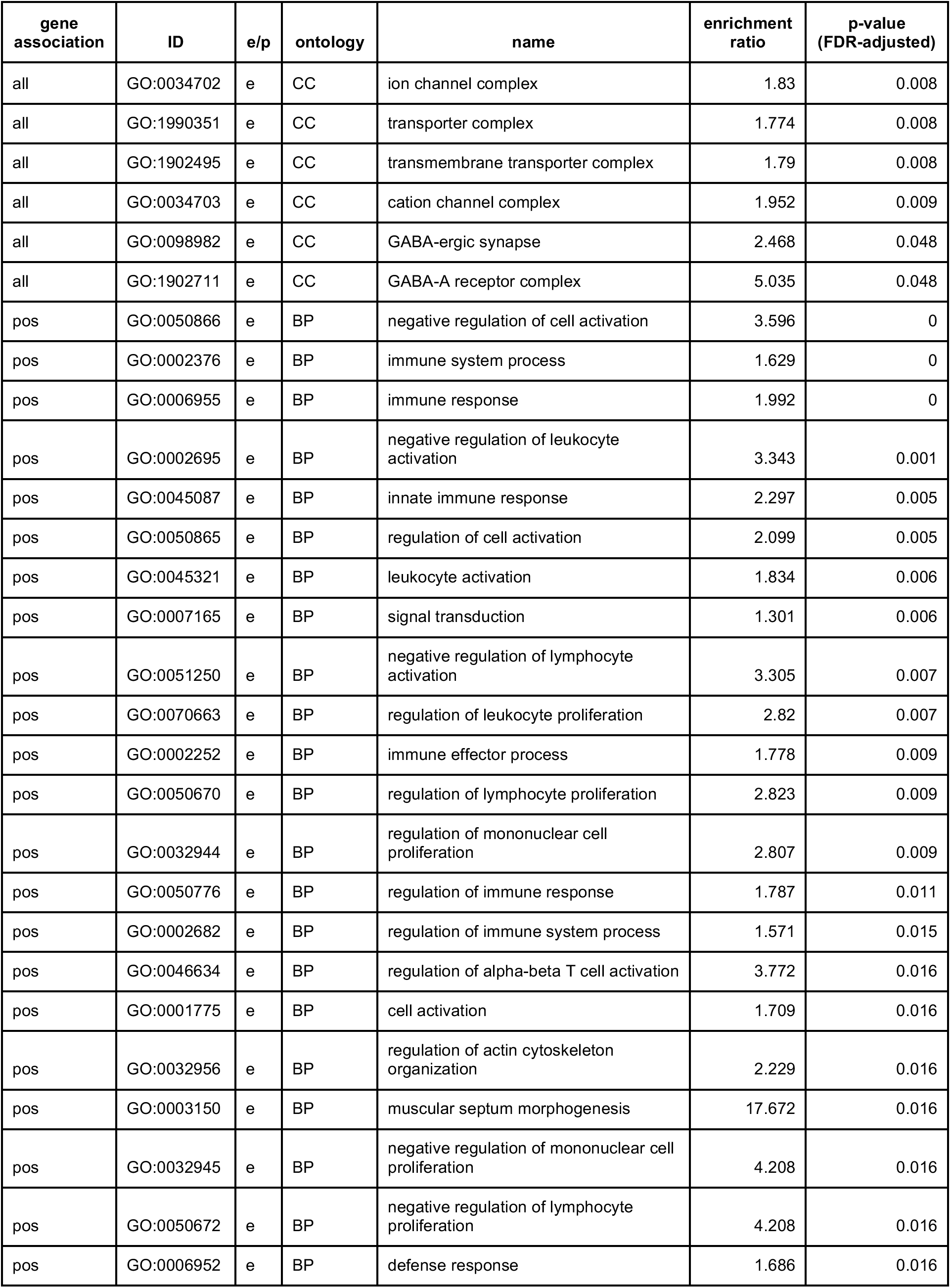
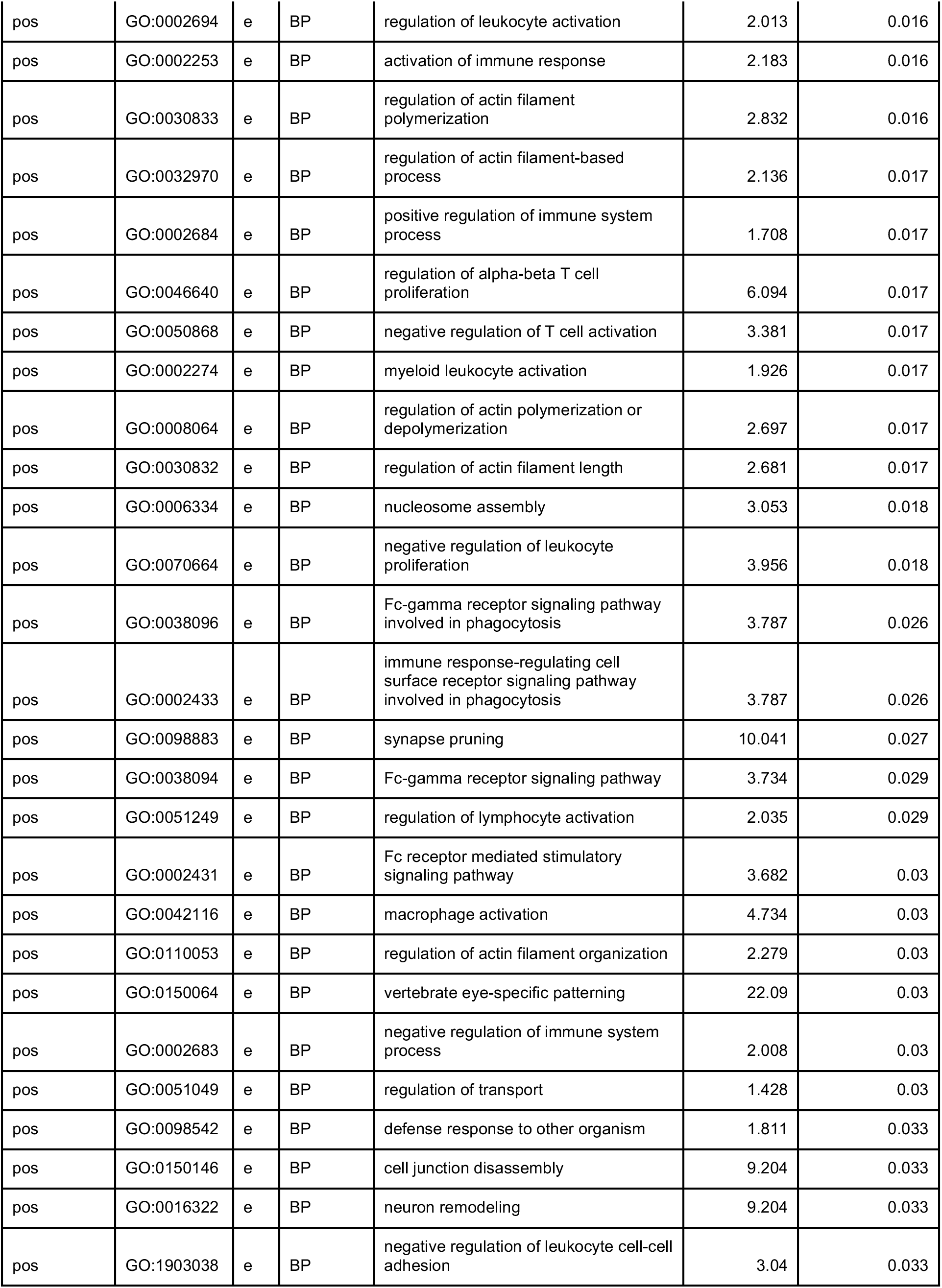
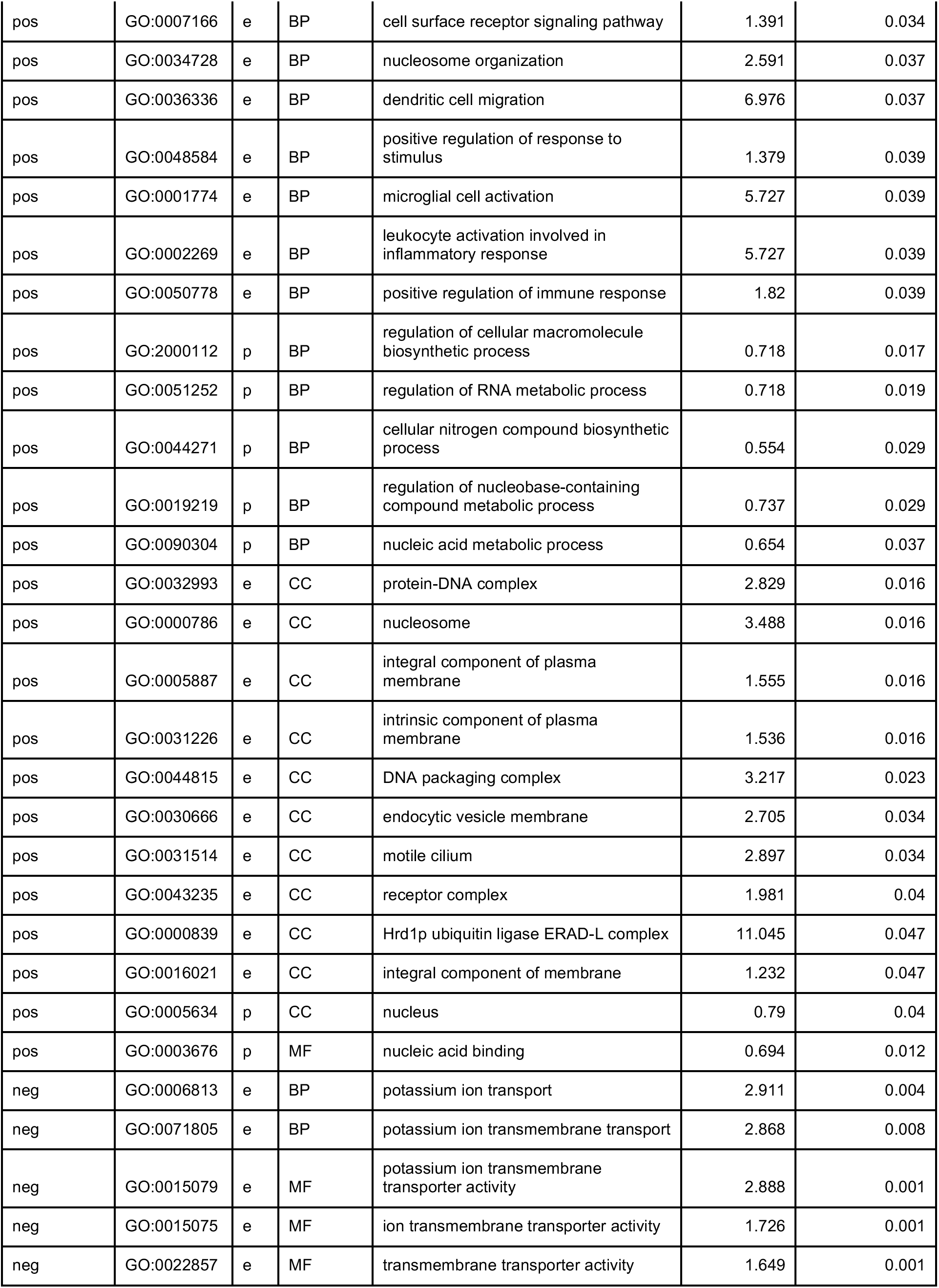
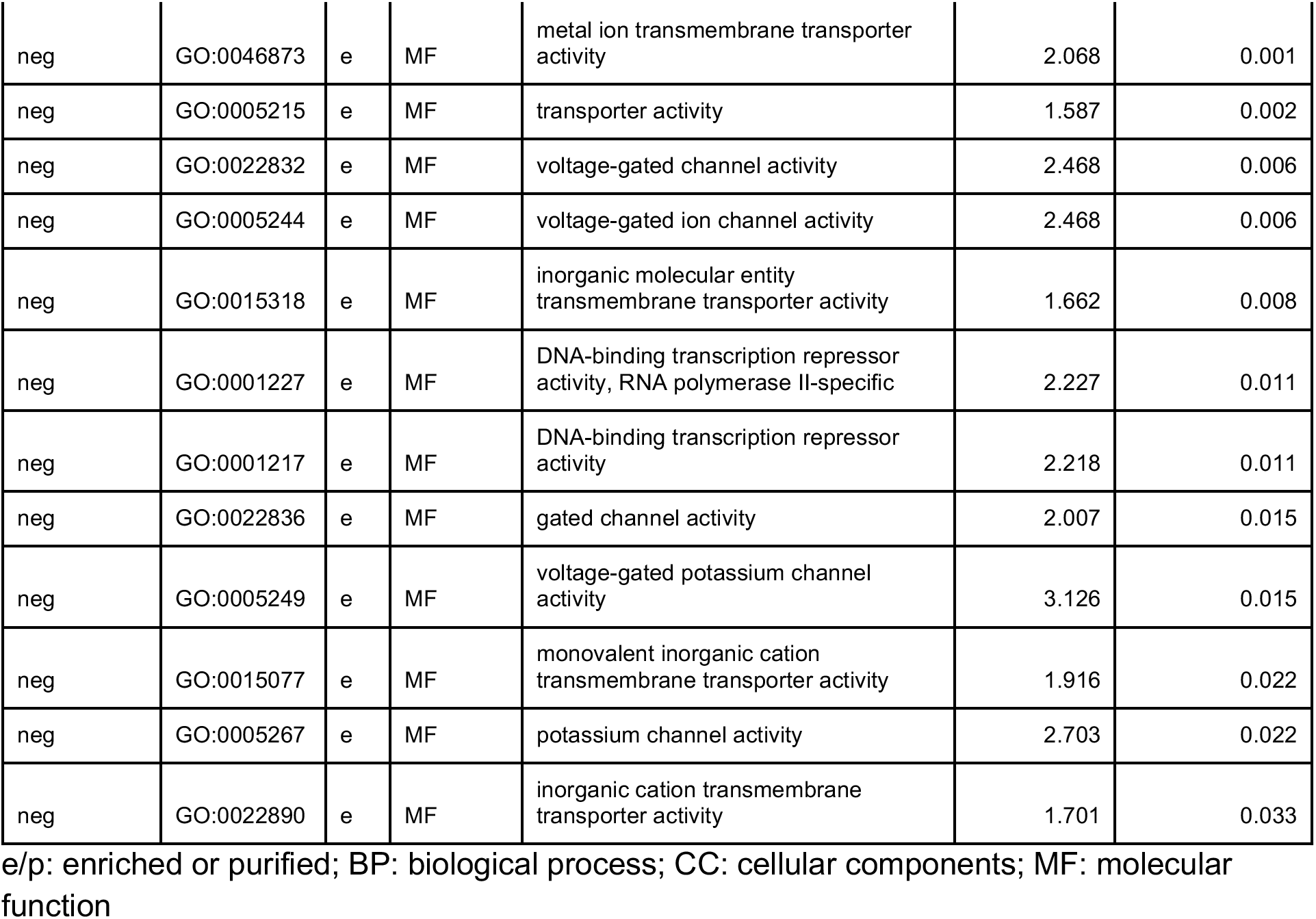
Significant items from all-gene GOEA

| gene association | ID | e/p | ontology | name | enrichment ratio | p-value (FDR-adjusted) |
| --- | --- | --- | --- | --- | --- | --- |
| all | GO:0034702 | e | CC | ion channel complex | 1.83 | 0.008 |
| all | GO:1990351 | e | CC | transporter complex | 1.774 | 0.008 |
| all | GO:1902495 | e | CC | transmembrane transporter complex | 1.79 | 0.008 |
| all | GO:0034703 | e | CC | cation channel complex | 1.952 | 0.009 |
| all | GO:0098982 | e | CC | GABA-ergic synapse | 2.468 | 0.048 |
| all | GO:1902711 | e | CC | GABA-A receptor complex | 5.035 | 0.048 |
| pos | GO:0050866 | e | BP | negative regulation of cell activation | 3.596 | 0 |
| pos | GO:0002376 | e | BP | immune system process | 1.629 | 0 |
| pos | GO:0006955 | e | BP | immune response | 1.992 | 0 |
| pos | GO:0002695 | e | BP | negative regulation of leukocyte activation | 3.343 | 0.001 |
| pos | GO:0045087 | e | BP | innate immune response | 2.297 | 0.005 |
| pos | GO:0050865 | e | BP | regulation of cell activation | 2.099 | 0.005 |
| pos | GO:0045321 | e | BP | leukocyte activation | 1.834 | 0.006 |
| pos | GO:0007165 | e | BP | signal transduction | 1.301 | 0.006 |
| pos | GO:0051250 | e | BP | negative regulation of lymphocyte activation | 3.305 | 0.007 |
| pos | GO:0070663 | e | BP | regulation of leukocyte proliferation | 2.82 | 0.007 |
| pos | GO:0002252 | e | BP | immune effector process | 1.778 | 0.009 |
| pos | GO:0050670 | e | BP | regulation of lymphocyte proliferation | 2.823 | 0.009 |
| pos | GO:0032944 | e | BP | regulation of mononuclear cell proliferation | 2.807 | 0.009 |
| pos | GO:0050776 | e | BP | regulation of immune response | 1.787 | 0.011 |
| pos | GO:0002682 | e | BP | regulation of immune system process | 1.571 | 0.015 |
| pos | GO:0046634 | e | BP | regulation of alpha-beta T cell activation | 3.772 | 0.016 |
| pos | GO:0001775 | e | BP | cell activation | 1.709 | 0.016 |
| pos | GO:0032956 | e | BP | regulation of actin cytoskeleton organization | 2.229 | 0.016 |
| pos | GO:0003150 | e | BP | muscular septum morphogenesis | 17.672 | 0.016 |
| pos | GO:0032945 | e | BP | negative regulation of mononuclear cell proliferation | 4.208 | 0.016 |
| pos | GO:0050672 | e | BP | negative regulation of lymphocyte proliferation | 4.208 | 0.016 |
| pos | GO:0006952 | e | BP | defense response | 1.686 | 0.016 |
| pos | GO:0002694 | e | BP | regulation of leukocyte activation | 2.013 | 0.016 |
| pos | GO:0002253 | e | BP | activation of immune response | 2.183 | 0.016 |
| pos | GO:0030833 | e | BP | regulation of actin filament polymerization | 2.832 | 0.016 |
| pos | GO:0032970 | e | BP | regulation of actin filament-based process | 2.136 | 0.017 |
| pos | GO:0002684 | e | BP | positive regulation of immune system process | 1.708 | 0.017 |
| pos | GO:0046640 | e | BP | regulation of alpha-beta T cell proliferation | 6.094 | 0.017 |
| pos | GO:0050868 | e | BP | negative regulation of T cell activation | 3.381 | 0.017 |
| pos | GO:0002274 | e | BP | myeloid leukocyte activation | 1.926 | 0.017 |
| pos | GO:0008064 | e | BP | regulation of actin polymerization or depolymerization | 2.697 | 0.017 |
| pos | GO:0030832 | e | BP | regulation of actin filament length | 2.681 | 0.017 |
| pos | GO:0006334 | e | BP | nucleosome assembly | 3.053 | 0.018 |
| pos | GO:0070664 | e | BP | negative regulation of leukocyte proliferation | 3.956 | 0.018 |
| pos | GO:0038096 | e | BP | Fc-gamma receptor signaling pathway involved in phagocytosis | 3.787 | 0.026 |
| pos | GO:0002433 | e | BP | immune response-regulating cell surface receptor signaling pathway involved in phagocytosis | 3.787 | 0.026 |
| pos | GO:0098883 | e | BP | synapse pruning | 10.041 | 0.027 |
| pos | GO:0038094 | e | BP | Fc-gamma receptor signaling pathway | 3.734 | 0.029 |
| pos | GO:0051249 | e | BP | regulation of lymphocyte activation | 2.035 | 0.029 |
| pos | GO:0002431 | e | BP | Fc receptor mediated stimulatory signaling pathway | 3.682 | 0.03 |
| pos | GO:0042116 | e | BP | macrophage activation | 4.734 | 0.03 |
| pos | GO:0110053 | e | BP | regulation of actin filament organization | 2.279 | 0.03 |
| pos | GO:0150064 | e | BP | vertebrate eye-specific patterning | 22.09 | 0.03 |
| pos | GO:0002683 | e | BP | negative regulation of immune system process | 2.008 | 0.03 |
| pos | GO:0051049 | e | BP | regulation of transport | 1.428 | 0.03 |
| pos | GO:0098542 | e | BP | defense response to other organism | 1.811 | 0.033 |
| pos | GO:0150146 | e | BP | cell junction disassembly | 9.204 | 0.033 |
| pos | GO:0016322 | e | BP | neuron remodeling | 9.204 | 0.033 |
| pos | GO:1903038 | e | BP | negative regulation of leukocyte cell-cell adhesion | 3.04 | 0.033 |
| pos | GO:0007166 | e | BP | cell surface receptor signaling pathway | 1.391 | 0.034 |
| pos | GO:0034728 | e | BP | nucleosome organization | 2.591 | 0.037 |
| pos | GO:0036336 | e | BP | dendritic cell migration | 6.976 | 0.037 |
| pos | GO:0048584 | e | BP | positive regulation of response to stimulus | 1.379 | 0.039 |
| pos | GO:0001774 | e | BP | microglial cell activation | 5.727 | 0.039 |
| pos | GO:0002269 | e | BP | leukocyte activation involved in inflammatory response | 5.727 | 0.039 |
| pos | GO:0050778 | e | BP | positive regulation of immune response | 1.82 | 0.039 |
| pos | GO:2000112 | p | BP | regulation of cellular macromolecule biosynthetic process | 0.718 | 0.017 |
| pos | GO:0051252 | p | BP | regulation of RNA metabolic process | 0.718 | 0.019 |
| pos | GO:0044271 | p | BP | cellular nitrogen compound biosynthetic process | 0.554 | 0.029 |
| pos | GO:0019219 | p | BP | regulation of nucleobase-containing compound metabolic process | 0.737 | 0.029 |
| pos | GO:0090304 | p | BP | nucleic acid metabolic process | 0.654 | 0.037 |
| pos | GO:0032993 | e | CC | protein-DNA complex | 2.829 | 0.016 |
| pos | GO:0000786 | e | CC | nucleosome | 3.488 | 0.016 |
| pos | GO:0005887 | e | CC | integral component of plasma membrane | 1.555 | 0.016 |
| pos | GO:0031226 | e | CC | intrinsic component of plasma membrane | 1.536 | 0.016 |
| pos | GO:0044815 | e | CC | DNA packaging complex | 3.217 | 0.023 |
| pos | GO:0030666 | e | CC | endocytic vesicle membrane | 2.705 | 0.034 |
| pos | GO:0031514 | e | CC | motile cilium | 2.897 | 0.034 |
| pos | GO:0043235 | e | CC | receptor complex | 1.981 | 0.04 |
| pos | GO:0000839 | e | CC | Hrd1p ubiquitin ligase ERAD-L complex | 11.045 | 0.047 |
| pos | GO:0016021 | e | CC | integral component of membrane | 1.232 | 0.047 |
| pos | GO:0005634 | p | CC | nucleus | 0.79 | 0.04 |
| pos | GO:0003676 | p | MF | nucleic acid binding | 0.694 | 0.012 |
| neg | GO:0006813 | e | BP | potassium ion transport | 2.911 | 0.004 |
| neg | GO:0071805 | e | BP | potassium ion transmembrane transport | 2.868 | 0.008 |
| neg | GO:0015079 | e | MF | potassium ion transmembrane transporter activity | 2.888 | 0.001 |
| neg | GO:0015075 | e | MF | ion transmembrane transporter activity | 1.726 | 0.001 |
| neg | GO:0022857 | e | MF | transmembrane transporter activity | 1.649 | 0.001 |
| neg | GO:0046873 | e | MF | metal ion transmembrane transporter activity | 2.068 | 0.001 |
| neg | GO:0005215 | e | MF | transporter activity | 1.587 | 0.002 |
| neg | GO:0022832 | e | MF | voltage-gated channel activity | 2.468 | 0.006 |
| neg | GO:0005244 | e | MF | voltage-gated ion channel activity | 2.468 | 0.006 |
| neg | GO:0015318 | e | MF | inorganic molecular entity transmembrane transporter activity | 1.662 | 0.008 |
| neg | GO:0001227 | e | MF | DNA-binding transcription repressor activity, RNA polymerase II-specific | 2.227 | 0.011 |
| neg | GO:0001217 | e | MF | DNA-binding transcription repressor activity | 2.218 | 0.011 |
| neg | GO:0022836 | e | MF | gated channel activity | 2.007 | 0.015 |
| neg | GO:0005249 | e | MF | voltage-gated potassium channel activity | 3.126 | 0.015 |
| neg | GO:0015077 | e | MF | monovalent inorganic cation transmembrane transporter activity | 1.916 | 0.022 |
| neg | GO:0005267 | e | MF | potassium channel activity | 2.703 | 0.022 |
| neg | GO:0022890 | e | MF | inorganic cation transmembrane transporter activity | 1.701 | 0.033 |

e/p: enriched or purified; BP: biological process; CC: cellular components; MF: molecular function

## Notes

https://github.com/rdgao/field-echos

### Competing Interest Statement

The authors have declared no competing interest.

### Summary of Updates

corrected various errors of figure referencing in main text, revised abstracted, and updated formatting

